# Early life imprints the hierarchy of T cell clone sizes

**DOI:** 10.1101/2020.07.21.214841

**Authors:** Mario U. Gaimann, Maximilian Nguyen, Jonathan Desponds, Andreas Mayer

## Abstract

The adaptive immune system responds to pathogens by selecting clones of cells with specific receptors. While clonal selection in response to particular antigens has been studied in detail, it is unknown how a lifetime of exposures to many antigens collectively shape the immune repertoire. Here, through mathematical modeling and statistical analyses of T cell receptor sequencing data we demonstrate that clonal expansions during a perinatal time window leave a long-lasting imprint on the human T cell repertoire. We demonstrate how the empirical scaling law relating the rank of the largest clones to their size can emerge from clonal growth during repertoire formation. We statistically identify early founded clones and find that they are indeed highly enriched among the largest clones. This enrichment persists even after decades of human aging, in a way that is quantitatively predicted by a model of fluctuating clonal selection. Our work presents a quantitative theory of human T cell dynamics compatible with the statistical laws of repertoire organization and provides a mechanism for how early clonal dynamics imprint the hierarchy of T cell clone sizes with implications for pathogen defense and autoimmunity.

## I. INTRODUCTION

The hallmark of adaptive immunity is the generation of diversity through genetic recombination and clonal selection. Their interplay balances the breadth and specificity of the ~10^12^ T cells in the human body (Fig. 1A) [1, 2]: The genetic recombination of the T cell receptor (TCR) locus, termed VDJ recombination, generates an enormous potential diversity of receptors ranging from early estimates of ~10^15^ [3] to more recent estimates of ~10^61^ [4] different possible receptor TCR*αβ* heterodimers. Clonal selection expands the number of specific cells during an infection for effector functions, a fraction of which are retained over prolonged periods of time as immune memory [2, 5].

**FIG. 1:**
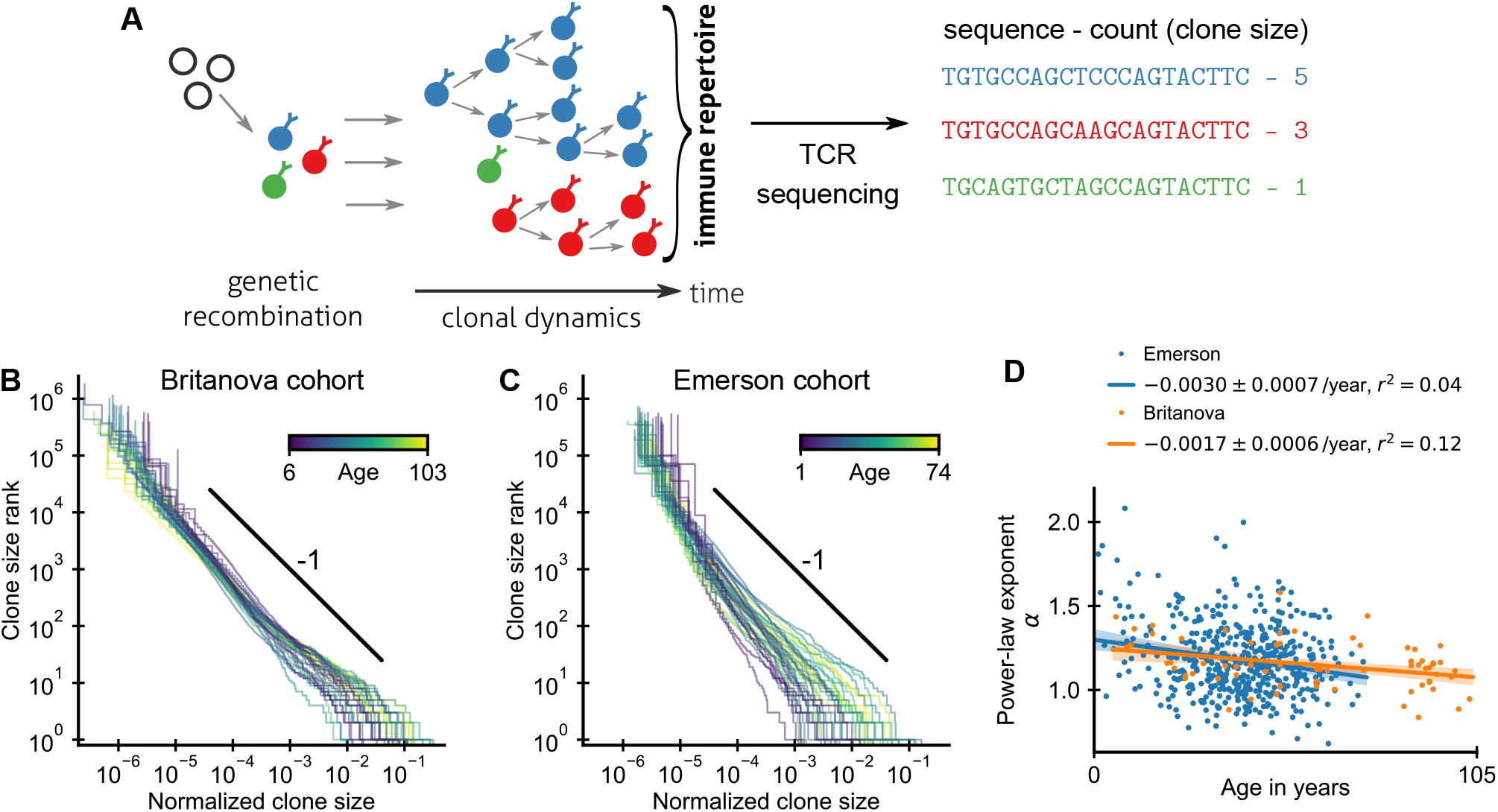
Statistics of human T cell repertoire organization. (A) T cells with highly diverse receptors are created from progenitor cells through genetic recombination (left), which then undergo clonal selection (middle) together shaping the immune repertoire. The T cell receptor (TCR) locus acts as a natural barcode for clonal lineages, which can be read out by sequencing (right). (B, C) Clone size distributions in two large cohort studies of human blood samples using disparate sequencing protocols display a power-law relationship between the rank and size of the largest clones. Each line shows the clone size distribution in an individual. Ages are color coded as indicated in the legend. The black line shows a power law with a slope of −1 for visual comparison. Clone sizes were normalized by the total number of reads and by the memory cell fraction to account for variations in sampling depth and in the subset composition of peripheral blood, respectively (Fig. S2). Only a single individual is displayed per two-year age bracket to improve visibility. (D) Power-law exponents as a function of the age (legend: linear regression slope and coefficient of determination). Data sources: B, D [11], C, D [12].

Much progress has been made deciphering the mechanisms of regulation and control of T cell dynamics over the last decades [2, 6, 7]. However, much of that progress has focused on the dynamics of subsets of T cells specific to a particular antigen and has come from experiments in mice. An important open question is how exposures to many antigens over a human lifetime collectively shape our T cell repertoire [2, 8].

High-throughput repertoire sequencing enables direct surveys of the diversity and clonal composition of T cells from human blood or tissue samples and thus promises to provide quantitative answers to this question [9–18]. However, while the TCR locus provides a natural barcode for clonal lineages due to its large diversity, this same diversity also makes inferring past clonal dynamics a challenging inverse problem, in particular given practical limitations on sequencing depth and temporal resolution in longitudinal studies. Mathematical modeling can help address this challenge by solving the forward problem of linking clonal dynamics to emergent statistical patterns [19–23]. Comparing patterns to data can provide insights about dynamics from static snapshots of repertoire organization in different individuals. A particularly striking such pattern has been the observation of power-law scaling of clone sizes spanning several orders of magnitude [9–13, 19]. In a typical sample of T cells from peripheral blood a large fraction (more than half in some individuals) of clones are only seen once within 10^5^ – 10^7^ sampled sequences. At the same time the most abundant clones typically account for more than 1% of all sequencing reads, equivalent to a clone size of ~10^10^ cells when extrapolating to the full repertoire. It is unknown when these large clonal expansions happen, and more broadly what determines the hierarchy of clone sizes.

Here, we use cohort and longitudinal human TCR repertoire sequencing data [11, 12, 17, 24] to develop a statistical theory of T cell dynamics. We find that clonal expansions during repertoire formation establish clone size scaling, and we show that clonal selection pressures during adult life only slowly reshape the initial hierarchy.

## II. RESULTS

### A. A scaling law of human T cell repertoire organization

An important statistic to summarize repertoire organization is the clone size distribution, which tabulates the number of clones found at different multiplicities within a repertoire or sample. Multiple previous studies have shown that these distributions are heavy-tailed [9–13], but potential confounding by noise introduced during the sequencing process has remain debated [22] and systematic analyses of how variable these distributions are across healthy individuals have been lacking. To fill these gaps we reanalyzed data from two large-scale cohort repertoire sequencing studies, which used fundamentally different sequencing pipelines and thus have different sources of noise (Material and Methods). Both studies sequenced the locus coding for the hypervariable TCR CDR3–*β* chain from peripheral blood samples of healthy human volunteers spanning a large range of ages (Fig. S1).

After normalizing clone sizes to account for variations in sampling depth and subset composition (Fig. S2), we found that the tails of the clone size distributions collapsed to the same statistical law across individuals and cohorts (Fig. 1B,C): Ranking clones by decreasing size, the rank of the largest clones approximately scales with their size *C* as a power law,

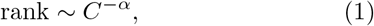

where *α* is a scaling exponent. To quantify the apparent similarity of the scaling relationship we determined *α* for each sample by maximum likelihood estimation. Only a small fraction of all T cells are sampled, which poses a challenge because subsampling a power law leads to deviations from scaling at small clone sizes [25]. To overcome this challenge we used a trimming procedure and excluded clones smaller than a minimal size from the fitting, which decreases bias arising from subsampling (SI Text C). Determined in this subsampling-robust manner the fitted power-law exponents agree remarkably well within the range of ages covered by both cohorts (Fig. 1D); with α = 1.17 ± 0.03 (mean ± standard error (SE)) and *α* = 1.18 ± 0.01 in the Britanova and Emerson cohort, respectively. Moreover, the fitted exponents varied little between individuals in both cohorts; with a sample standard deviation of fitted exponents of 0.14 and 0.21, respectively. The agreement of the mean exponents is noteworthy given the different sequencing pipelines and provides strong evidence that the scaling relationship (Eq. 1) is a true feature of the clone size distribution and not of the measurement process.

What drives the emergence of a power-law distributed hierarchy of clone sizes? Given the reproducibility of the scaling law across individuals we might hope for a statistical explanation independent of the precise antigenic history that drives the expansion of specific cells in an individual. To test hypotheses about mechanisms underlying scaling we describe repertoire dynamics using a general mathematical framework based on effective stochastic rate equations for the recruitment of new clones, and the proliferation and death of already existing clones within a T cell compartment (Material and Methods). In macroecology, where such reductionist approaches have a long history, simple neutral models within this framework have had surprising success in describing species abundance distributions only accounting for demographic stochasticity [26], but this source of variability is insufficient to account for the observed breadth of T cell clone sizes [19, 23] (for a detailed discussion see SI Text E). The failure of this null model has prompted a search for other mechanisms that explain scaling.

To constrain this search we analyzed how fitted exponents varied with age. In particular, we expected a substantially steeper tail in young individuals based on a finite time solution we derived for a previously proposed model of how power-law scaling can emerge from the cumulative effect of temporal fluctuations in clonal growth rates [19] (SI Text G 1). While exponents overall decreased slightly with age, the dependence on age accounted for surprisingly little variation in both cohorts (Fig. 1D and Fig. S3). Notably, scaling is established within the first decade of life, with significant clone size variability existing as early as at birth (Fig. S4), defying previous model predictions.

### B. A mechanism for the emergence of scaling during repertoire formation

We hypothesized that scaling might result from clonal expansions during repertoire formation, which would naturally explain the early onset of scaling. Our hypothesis is based on experimental evidence in mice [27, 28] and human [29, 30] that repertoire formation is driven not only by increased thymic output, but also by large proliferative expansion of some T cell clones. Additionally, multiple studies [31–33] have shown that some T cell clones can persist over multiple decades, which suggested to us that clonal turnover might be sufficiently slow (see also SI Text E 2) for transient expansionary dynamics early in life to shape repertoire organization over prolonged periods of time.

To test our hypothesis we constructed a minimal model of repertoire formation based on known T cell biology (Fig. 2A). Following previous work [20, 34] we assume that the proliferation rate *b* is inversely proportional to the total number of cells already in the repertoire to model increased proliferation early in life. This dependence of proliferation rate on repertoire size arises in a simple mechanistic model of T cell competition (SI Text F 1). For simplicity we further assume that the rates of cellular death *d* and recruitment of new clones *θ* are constant. Importantly, recruitment of new clones and total expansion of already existing clones maintain a constant ratio throughout development under these assumptions in line with findings that the fraction of cells with T cell receptor excision circles, which are diluted during peripheral division, is constant during fetal development [30] and infancy [34].

**FIG. 2:**
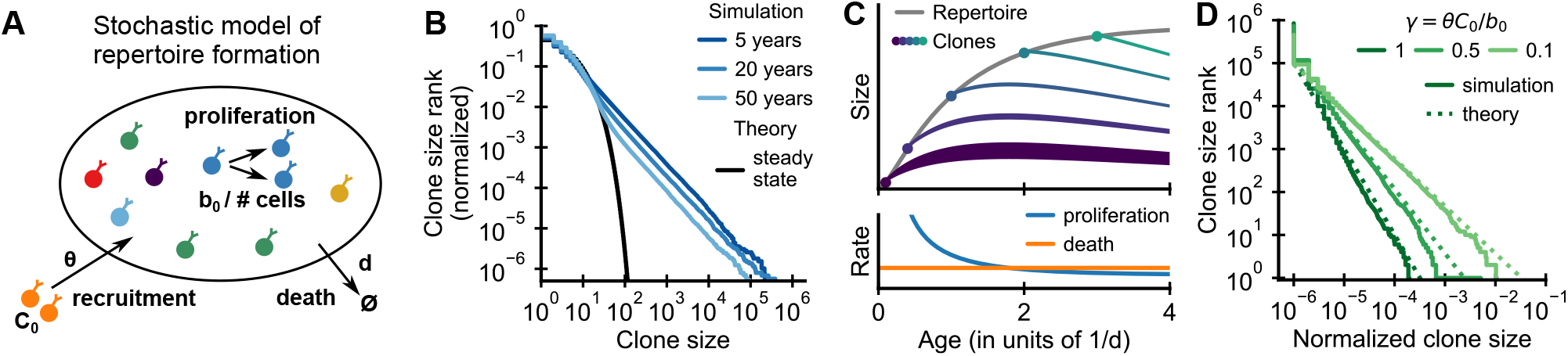
Emergence of power-law scaling of clone sizes in a minimal model of repertoire formation. (A) Sketch of the stochastic dynamics of recruitment, proliferation, and death of T cells. Proliferation is inversely proportional to total repertoire size modeling increasing competition during repertoire growth. (B) Clone size distributions in simulated repertoires display power-law scaling (blue lines), in contrast to steady-state predictions (black line, SI Text Eq. 14). (C) Illustration of the mechanism: Early in life rates of proliferation exceed clonal turnover (lower panel). As the total repertoire size increases (grey line, upper panel) the proliferation rate decreases due to increased competition. The dynamics of selected clones after their recruitment marked by a dot is indicated by colored lines (upper panel). The line position shows the cumulative size of all prior clones, while the line width indicates the size of the clone (not to scale). The earlier a clones is recruited the larger it expands during the period of overall repertoire growth. (D) Dependence of the clone size distribution on parameters. Simulated repertoires at 5 years of age were subsampled to 10^6^ cells to mimick the experimental sampling depth (solid lines). The simulated data closely follow predictions from a continuum theory of repertoire formation (dashed lines). Parameters: (B,D) *d* = 0.2/year, *C*_0_ = 1, *θ* = 10^6^/year; (B) *γ* = 0.1 (implying *b*_0_ = 10^7^/year).

We simulated the model starting from an empty repertoire and found that large clones displayed power-law scaling (Fig. 2B blue lines). The simulation results contrast with steady state predictions (Fig. 2B black line), where the model effectively reduces to the neutral null model introduced earlier (SI Text F4). Thus we find that repertoire formation can produce transient but long-lasting power-law scaling of clone sizes.

To obtain intuitive insight into how scaling is established, we developed a continuum theory of clonal dynamics during repertoire growth (SI Text F 3). We find that the clone size *C_i_* of the *i*-th clone recruited at time *t_i_* follows a subexponential growth law *C_i_*(*t*) = *C*_0_ (*t*/*t_i_*)^1/(1+*γ*)^, where *γ* is the ratio of the contribution of recruitment and proliferation to overall compartment growth. Clones recruited early grow large deterministically until competition lowers proliferation rates below the death rate (Fig. 2C, lower panel). Different clones are recruited at different times and thus have more or less time to grow (Fig. 2C, upper panel), which leads to a clone size distribution that follows power-law scaling with an exponent *α* = 1 + *γ*. We note that this origin of the power-law scaling is closely related to a well-known generative mechanism for power-laws first studied by Yule [35] (for a detailed discussion see SI Text F6).

The predicted exponent closely matches simulation results for different values of *γ* (Fig. 2D dashed lines). Intuitively, when recruitment rates are higher clones founded early have less time to outgrow later competitors, and thus the power law is steeper (*α* is larger). Importantly, in the biological parameter regime in which proliferation dominates, *γ* < 1, the exponent is compatible with experiments (Fig. 1B-E). We thus find, that the model – without fine tuning of parameters – reproduces the observed scaling exponent.

To expose a basic mechanism capable of producing broad clone size distributions we have kept the model deliberately simple. More detailed models demonstrate the conditions and limits on the generalizability of this mechanism (SI Text F 5). Variable recruitment sizes only affect the distribution of small clones (SI Text S19); while a saturation of proliferation rates, or competition between subsets of T cells for specific resources maintain distributions at small and intermediate sizes while leading to cutoffs for the largest clones (Fig. S17 and Fig. S18).

### C. Long-lived incumbency advantage shows early expansions imprint clone size hierarchy

Our proposed theory for the rapid emergence of scaling predicts that large clones have expanded massively during repertoire formation. To test this prediction we need to trace the dynamics of early founded clones. To this end, we exploit a change in the recombination statistics taking place during fetal development [36–38] (Fig. 3A). While T cells are produced by the thymus from the late first trimester the enzyme terminal deoxynucleotidyl transferase (TdT), which inserts non-templated nucleotides during VDJ recombination, is not expressed until the mid second trimester [38]. Therefore many more T cells in fetal and neonatal blood have zero insertions than expected by the adult recombination statistics [37]. This enables a statistical dating of individual clones in a repertoire based on their sequence [32, 39].

**FIG. 3:**
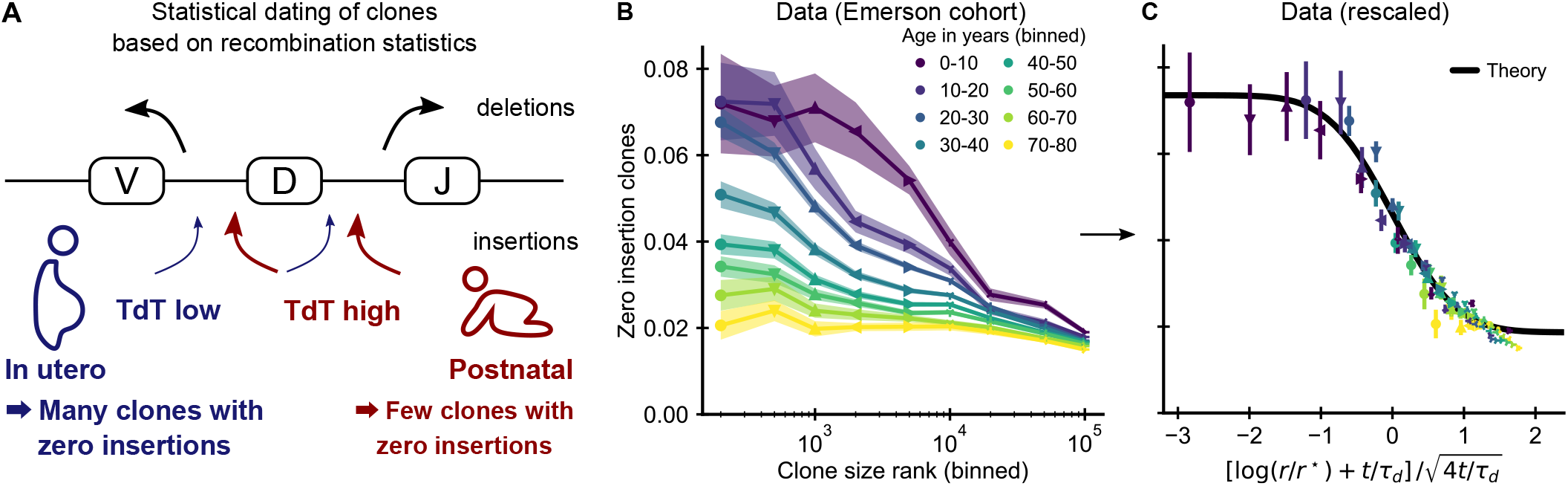
Statistical dating of clones reveals that early expansions have a long-lasting effect. (A) Genetic recombination of a TCR involves the choice of a V, D, and J region among multiple genomically-encoded templates as well as the deletion and insertion of nucleotides at both the VD and DJ junction. The enzyme TdT, which is responsible for nucleotide insertions, is not expressed during early fetal development. This allows a statistical dating of clonal ages, as clones with zero insertions at both junctions constitute a much larger fraction of all clones during a fetal and perinatal time window. (B) Fraction (± SE) of clones with zero insertions as a function of age and clone size. Clones are binned by their size into non-overlapping bins (rank 1 to 500, 501 to 1000, and so on; upper values are indicated on the x-axis). (C) Same data as in B displayed with a rescaled x-axis using fitted parameters *τ_d_* = 9.1 ± 0.5 years, *r*\* = 1.2 ± 0.2 · 10^4^. The data collapses onto a sigmoidal function predicted by theory (SI Text Eq. 51) with fitted *p*_0,−_= 0.074 ± 0.004, *p*_0,+_ = 0.0187 ± 0.0005 (black line). Data source: [12].

If our model is correct we expect abundant clones to be more likely to have zero insertions than smaller clones. Analyzing data from the Emerson cohort we find that zero insertion clones are indeed highly enriched within the most abundant clones (Fig. 3B). This generalizes a previous report of such an enrichment within the naive compartment [32]. The large cohort size allows us to perform a fine-grained analysis of how the fraction of zero-insertion clones depends on clonal abundance and age. We find that enrichment is particularly pronounced in the young and decreases with age at different speeds depending on clone size. Among the largest clones many more still have zero insertions than expected under the adult recombination statistics even multiple decades after repertoire formation. This suggests that the incumbent large clones created during repertoire formation are only slowly replaced by clones expanding later in life. Additional analyses rule out other potential explanations for the relation between insertion statistics and clonal abundance. Firstly, sequences with zero insertions are similarly enriched among the largest clones in productive and unproductive sequences (Fig. S5) demonstrating that convergent selection pressures during adult life are not a primary source of the higher abundance of these clones. Secondly, while abundant clones are also enriched for sequences with known antigen specificity (Fig. S6) and sequences likely to be convergently recombined (Fig. S7), these enrichment do not show the same striking dependence on age. Furthermore, we find that zero insertion clones were consistently less enriched in individuals infected by cytomegalovirus (Fig. S8), in contrast to the hypothesis that this infection might drive their expansion [32]. Taken together, these analyses support the conclusion that dynamics during the perinatal time window of repertoire formation leave a long-lasting imprint on the T cell clonal hierarchy well into adulthood.

### D. Longitudinal clone size fluctuations predict the dynamics of the clone size hierarchy with aging

Building on this successful validation of a core prediction of our theory we asked whether we could leverage the detailed pattern of enrichments at different ranks and ages to quantify how much being part of the wave of early expansions determines the fate of a clone relative to other sources of clone size variability. To this end we extended our model beyond repertoire formation and allowed clonal proliferation rates to fluctuate over time to model the net effect of clonal selection by changing antigenic stimuli during adult life [19] (Eq. 6).

To determine a biologically plausible fluctuation strength we analyzed the variability of clone sizes over time in a longitudinal study of T cell dynamics [24]. We first analyzed to what extent recently expanded clones contribute to the tail of the clone sizes, and found that only a small fraction of the largest clones in any sample were not already large at the earliest time point (Fig. 4A and Fig. S9). To minimize confounding by transient dynamics affecting these clones, we excluded these clones from further analysis. We found that large clones had remarkably stable abundances over time, which we quantified by calculating the variance of log-foldchanges in clone size between the second and every subsequent time point (Fig. 4B and Fig. S10). The variability of clone sizes increased linearly over time as expected theoretically, from which we determined a magnitude of net growth rate fluctuations compatible with the slope of increase (SI Text G3).

**FIG. 4:**
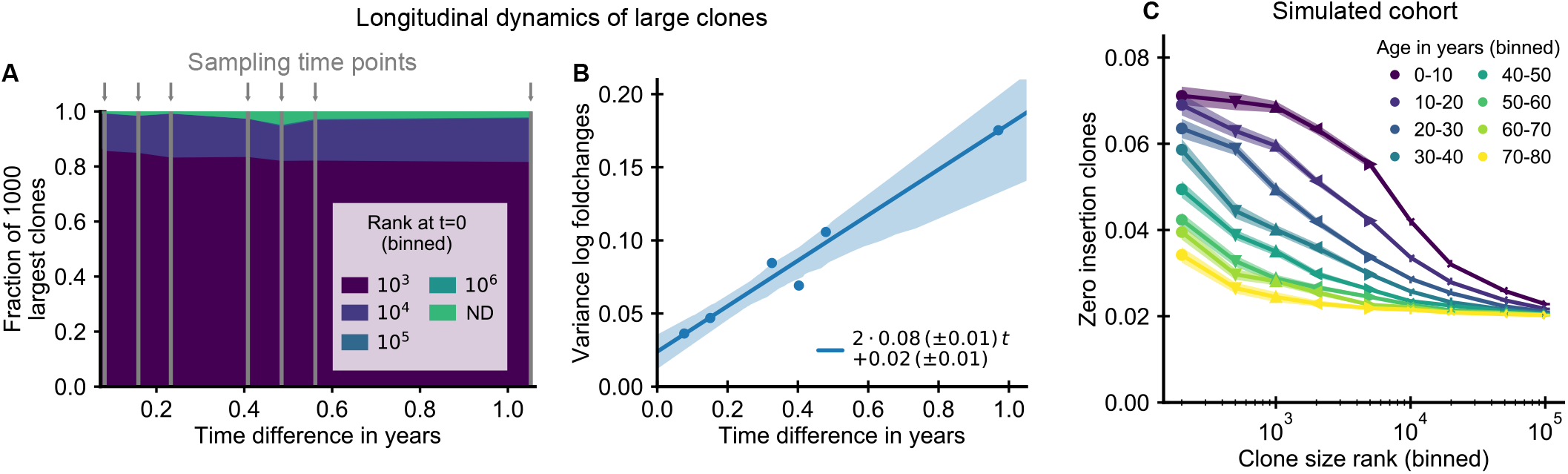
The small magnitude of longitudinal clone size fluctuations implies a slow reordering of the clone size hierarchy. (A,B) Longitudinal clonal dynamics in a healthy adult over a one year time span. (A) Fraction of the 1000 largest clones that fall within a specific clone size rank bin at the earliest time point. A small number of clones was not detected at all at the first time point (ND) likely representing recently expanded clones. All other clones were already among the largest clones initially. (B) Variance of log-foldchanges in clone size as a function of time difference for the 250 largest clones. (C) Fraction of clones with zero insertions as a function of age and clone size in a simulated cohort using a magnitude of clonal growth rate fluctuations inferred from the longitudinal data. Data source: [24].

Using the fitted fluctuation strength we constructed an *in silico* cohort of individuals of different ages according to the extended model (SI Text B3). In short, we computationally assigned each newly recruited clone to have zero insertions in a way that mimicks the change in fetal recombination statistics, and we simulated memory repertoire dynamics based on the combined effect of early expansion and fluctuating clonal selection. The enrichment of zero insertion clones in the simulated cohort (Fig. 4C) closely recapitulated the empirical findings using plausible parameter values (SI Text B4). Notably, the more long-lasting enrichment of zero insertion clones among the very largest clones is also found in the simulated cohort, and the timescales over which the enrichment decays agree remarkably well.

For a direct comparison between theory and experiment we mathematically analyzed how fluctuating selection reorders the initially established clone size hierarchy. The analytical results suggest a two-parameter rescaling of the enrichment of zero insertion clones as a general test of our theory (SI Text G 4). The two parameters of the theory, *τ_d_* and *r*\*, can be fitted from the enrichment data (SI Text B2). Rescaling the data with the fitted parameters leads to a collapse of all data points onto a single curve predicted by theory for both the simulated (Fig. S11) and experimental cohort (Fig. 3C). The fitted parameters quantify key features of long-term repertoire dynamics, with *τ_d_* characterizing the timescale over which fluctuations change the clone size hierarchy, and *r*\* being related to the number of clones recruited during early repertoire growth. In line with the long-lived enrichment of zero insertion clones, the fitting reveals a remarkably slow timescale of about a decade over which the clone size hierarchy is reordered during healthy aging. The fitted *r*\* indicates that early repertoire formation involves the expansion of a large number of different clones. Overall, the agreement between theory and data demonstrates that our model quantitatively captures how early expansions and ongoing fluctuating selection together shape the clone size hierarchy.

## III. DISCUSSION

The evolution of the adaptive immune system has endowed vertebrates with the ability to adapt to pathogens that evolve on a timescale faster than host reproduction [40]. However, this ability comes with a cost: every generation needs to rebuild immune memory anew. As the organism first comes into contact with the outside world it quickly needs to train its adaptive immune system to tolerate innocuous antigens and build up immune memory against pathogens. Here, we have shown that this process of rapid adaptation leaves a long-lasting imprint on the organization of the human T cell repertoire. More broadly, we propose a theory of repertoire dynamics that quantitatively describes how early expansions during repertoire formation combine with a lifetime of exposures to cumulatively shape the T cell hierarchy. Notably, we find that the T cell repertoire is remarkably stable over time in adult individuals outside of the punctuated expansions and contractions of specific clones in acute responses. Our study demonstrates that despite its vast complexity repertoire dynamics is partially predictable by quantitative models. The model predictions can help guide future longitudinal studies, which in turn will allow refinements of modeling assumptions. The current work thus provides a stepping stone towards a detailed quantitative understanding of T cell dynamics that we hope will ultimately power the rational development of immunodiagnostics and therapeutics.

The general mechanism we describe for imprinting in the adaptive immune system provides a unified lens through which to view a number of converging lines of evidence about how a developmental time window shapes adaptive immunity [2, 41–46]. In our model, overall repertoire growth early in life amplifies the effect of any early exposures, as they lead to much larger clonal expansions than similar exposures happening after the homeostatic repertoire size is reached. We thus expect early pathogen exposures to be particularly potent, as has been observed in influenza, where disease severity across age cohorts for different strains depends on the first exposure [42]. Conversely, we expect the presence of tolerizing factors early in life to be particularly crucial during repertoire formation to avoid autoimmunity, as has been observed for the autoimmune regulator gene AIRE, for which expression is only essential during a perinatal time window [41].

A limitation of datasets used in this study is that they do not provide direct information about the phenotypic characteristics of cells belonging to different clones. Repertoire sequencing of phenotypically sorted blood samples shows that the largest clones predominantly consist of cells with memory phenotype (SI Text D). This indirectly suggests that the clonal expansions during repertoire formation produce memory cells as we have assumed in our simulated cohort (Fig. 4C). Supporting this interpretation, a substantial number of memory cells circulate in the blood quickly following birth [47] and recent evidence suggests that memory-like T cells are already generated in the human intestine even before birth [44]. However alternatively, early expansions could also set up a broad distribution of naive T cell clone sizes [23], whose hierarchy would then need to be roughly maintained during the transition into memory to be compatible with the observed impact of early expansions on the hierarchy of the most abundant clones. Advances in single-cell technologies linking TCR sequencing and cellular phenotyping could help differentiate between these scenarios in the future.

An important questions raised by our work is which antigens drive the expansion of early T cell clones. To address this question it will be necessary to determine the exposures that imprint the abundance of these clones, as has been done recently for mucosal-associated invariant T cells [43], a subset of non-conventional T cells. Going forward, the highly abundant clones with sequences close to the genetically inherited gene templates resulting from the absence of TdT expression during early fetal development are a particularly interesting target of study. They might constitute an evolutionarily controlled set of innate-like defenses within the adaptive immune system. Determining what imprints their abundances will help resolve the question of whether their large abundances are simply a byproduct of rapid repertoire formation or whether these clones serve particular functions.

## IV. MATERIAL AND METHODS

### A. Repertoire sequencing data

We analyzed T cell repertoire sequencing data from the two largest published cohort studies of healthy human volunteers by Britanova *et al*. [11] and Emerson *et al*. [12] and from a longitudinal study by Chu et *al*. [24], detailed descriptions of which are provided in the SI Text Extended Methods.

In short, Britanova *et al*. [11] sequenced reverse transcribed mRNA with added unique molecular identifiers (UMIs), while Emerson *et al*. [12] and Chu *et al*. [24] sequenced genomic DNA coding for this region without the addition of UMIs. These approaches have complementary strengths: The addition of UMIs allows to correct for stochasticity during polymerase chain reaction (PCR) amplification and sequencing artifacts, while DNA sequencing removes the influence of cell-to-cell gene expression heterogeneity.

### B. Mathematical framework

We describe T cell dynamics using the following general set of stochastic rate equations. The class of models we consider are known in the mathematical literature as birth-death-immigration models. The number of cells *C_i_*, *i* = 1, …, *M* of each of the *M* clones in the repertoire changes according to

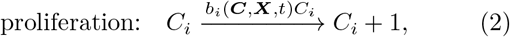

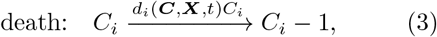

where the rate of proliferation *b_i_*(***C***, ***X***, *t*) or cell death *d_i_*(***C***, ***X***, *t*) generally can depend on the repertoire composition ***C***, on the time *t*, and on the state of the environment ***X***(*t*) representing e.g. the levels of different antigens and cytokines in the organism at a given time. We furthermore consider that new clones are added at rate *θ*(***X***, *t*) at a size *C*_0_,

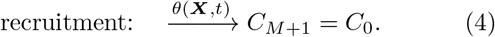

This recruitment represents thymic output and antigen-driven differentiation of naive cells for the naive and memory compartment, respectively.

In Sec. II B we study the influence of repertoire formation on clone sizes under the following assumptions:

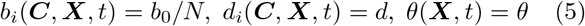

where 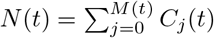 is the total repertoire size. In Sec. II C we modify this model by adding a noise term that describes the effective influence of environmental variations on clonal proliferation,

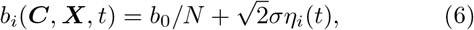

where 〈*η_i_*(*t*)*η_j_*(*t*′)〉 = *δ_ij_δ*(*t* – *t*′).

## Acknowledgements

We thank William Bialek, Curtis Callan, Ivana Cvijovic, Yuval Elhanati, Simone Mayer, Mikhail Pogorelyy, and Ned Wingreen for discussions and comments on the manuscript. This work was supported by a DAAD RISE Worldwide fellowship (MUG), the NSF-Simons Center for Quantitative Biology under grants Simons Foundation SFARI/597491-RWC and National Science Foundation 17764421 (JD), and a Lewis–Sigler fellowship (AM).

## Author contributions

AM conceptualized the problem and supervised research, MUG and AM wrote the draft manuscript, all authors performed analytic calculations, simulations, and reviewed the manuscript.

