## Supplemental Text and Figures for "Early life imprints the hierarchy of T cell clone sizes"

### Contents

**A. Supporting Figures** 2

**B. Extended Methods** 10
1. Data sources 10
2. Data analysis 10
3. Simulation procedures 11
4. Parameter choices 12

**C. Subsampling scaling** 12
1. Inference of scaling exponent 13
2. Graphical display of subsampled distributions 14

**D. Relation between clone size and cellular phenotypes** 15

**E. Modeling neutral repertoire dynamics** 17
1. Steady state clone size distribution 17
2. Relaxation time scale 17

**F. Modeling repertoire formation** 18
1. Mechanistic motivation for the competition function 18
2. Mean-field competition approximation 18
3. Continuum theory of clonal growth 18
4. Steady-state distribution 19
5. Relaxations of model assumptions 19
6. Relation to mechanisms generating power laws in other growth processes 22

**G. Modeling long-term repertoire dynamics with fluctuating clonal growth rates** 23
1. Slow convergence to steady-state scaling 23
2. A note on the scaling exponent 24
3. Predictions for longitudinal fluctuations in clone sizes 25
4. Relaxation of the zero insertion distribution 25

**References** 25

### A. SUPPORTING FIGURES

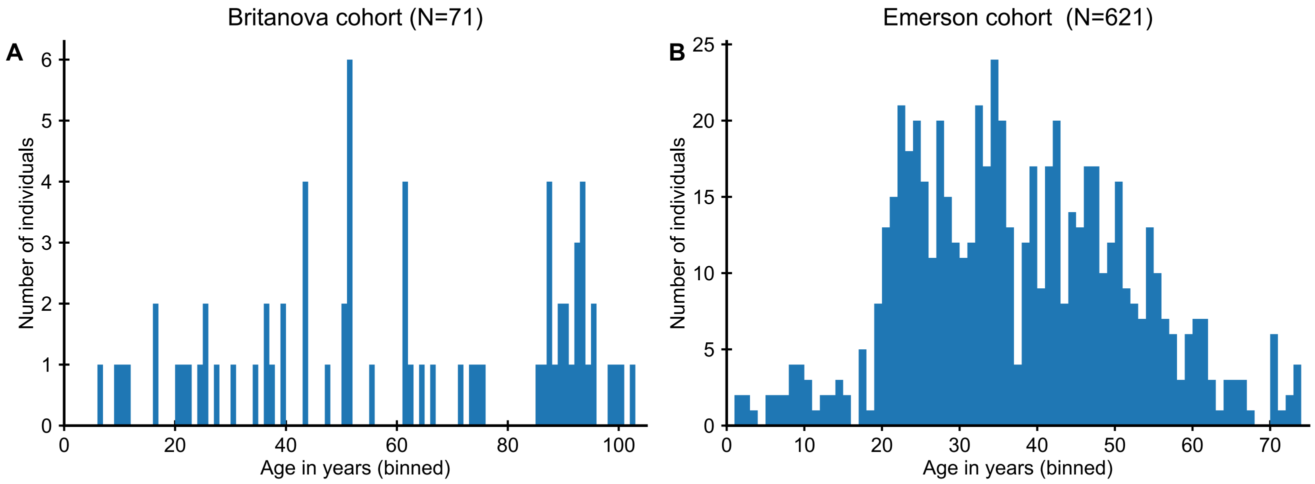


FIG. S1: **Distribution of ages in the two cohort studies.**

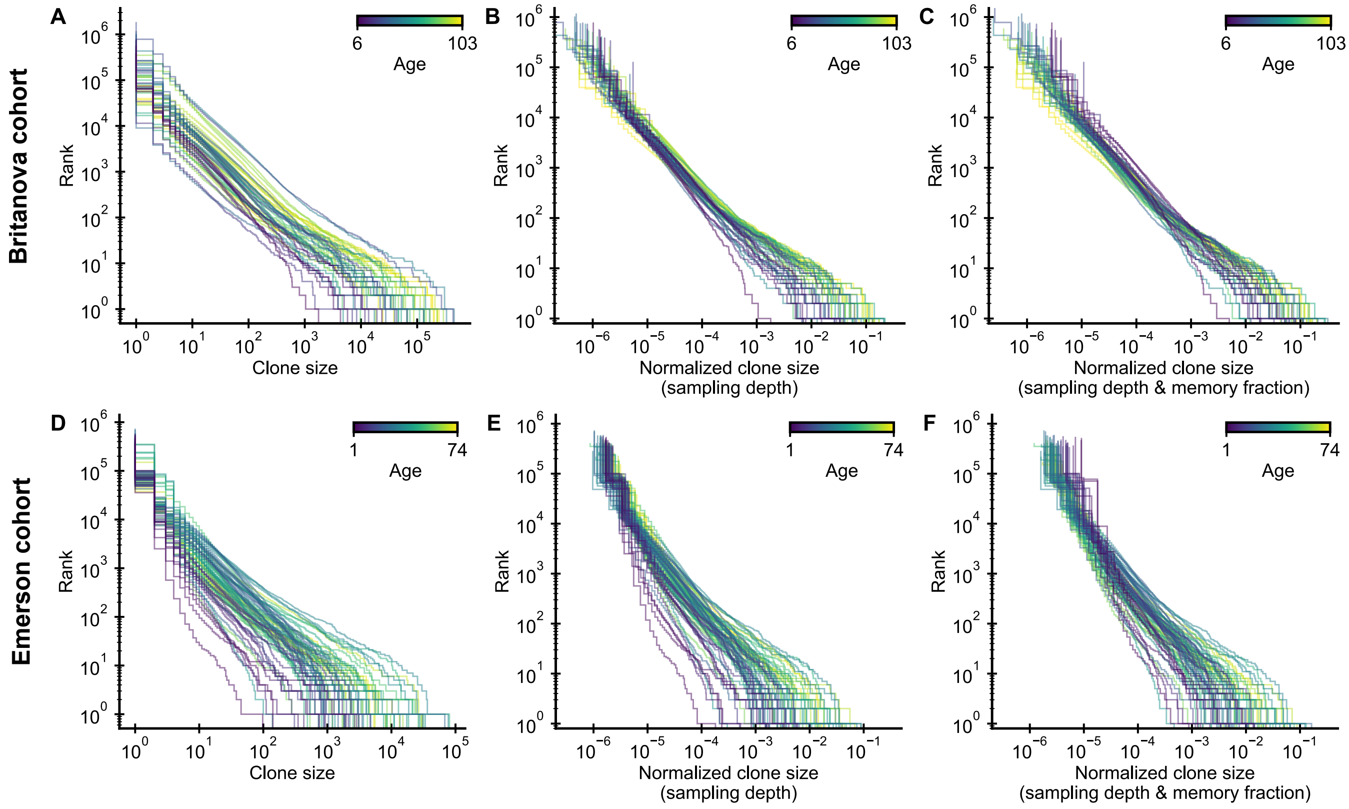


FIG. S2: **Influence of normalization choice on clone size distributions (see Extended Methods B 2).** (A,D) Raw clone size distributions show large variability due to different sample sizes. (B,E) A normalization by sampling depth removes much of this variation. (C,F) A normalization by the fraction of memory cells at different ages further collapses the tails of the clone size distributions. Data sources: A-C [1], D-F [2].

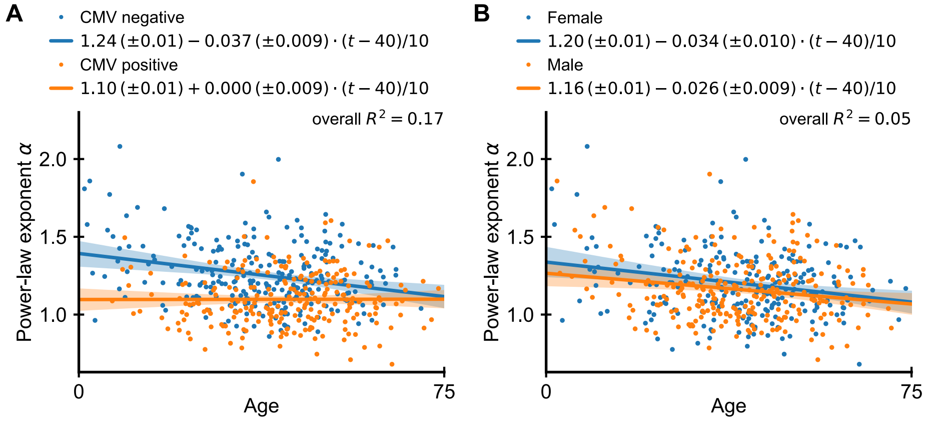


FIG. S3: **Dependence of power-law exponent on age by cytomegalovirus (CMV) infection status and sex.** (A) Chronic infection with CMV drives large clonal expansions [3, 4]. We thus repeated the analysis of Fig. 1E separating individuals based on their CMV infection status (fitted lines shown in legend, regression results displayed as offset + slope · (age in years - 40)/10). Overall, CMV positive individuals have a smaller $\alpha$ than uninfected individuals, which is independent of age. The average exponent in CMV negative individuals decreases slowly with age, and in old age coincides those of CMV positive individuals. Combining CMV infection status and age explained a significantly larger proportion of the variance in scaling exponents (17%) than age alone. (B) Many immune determinants differ markedly between the sexes [5]. We thus analyzed whether $\alpha$ depends on sex. We find that the dependence on age is similar among the sexes, but men have on average a slightly smaller exponent than women indicating a more skewed repertoire organization. Data source: Emerson *et al.* [2].

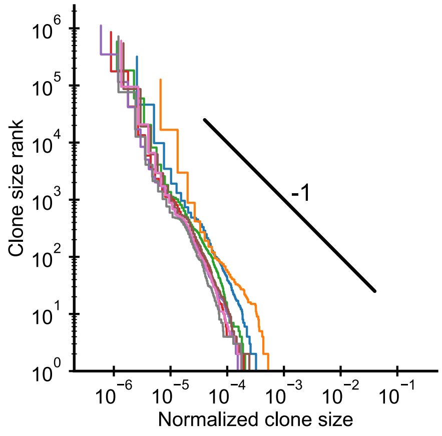


FIG. S4: **Clone size distributions of human T cell receptor repertoires in cordblood.** Each line shows the distribution in one individual. The black line shows a power law with a slope of -1 for visual comparison. The fitted power-law exponents $\alpha = 2.1 \pm 0.1$ (mean $\pm$ SE) are larger than in adult repertoires, but clone sizes are already remarkably broad. Data source: Britanova *et al.* [1].

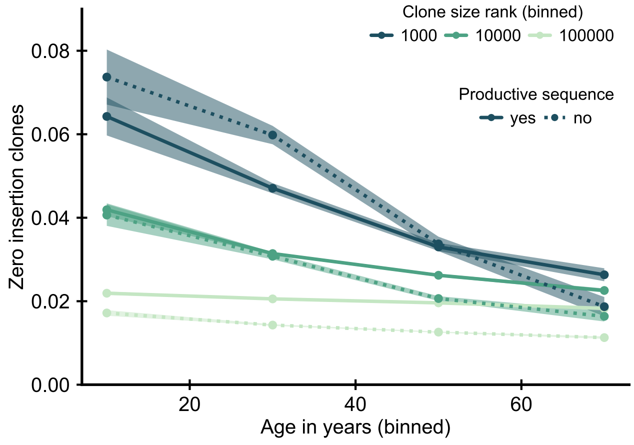


FIG. S5: **Comparison of the relative fraction of zero insertion clones within productive and unproductive sequences.** Sequences with zero insertions code for a particular subset of all possible TCRs, and some of their enrichment might represent a peripheral selective advantage of this subset of receptors. We thus asked how the enrichment depends on whether the sequence used to define the clone represents a productive or unproductive rearrangement. An unproductive rearrangement, in which the recombination process introduces a frameshift or stop codon, can be rescued by a second productive rearrangement, but is not expressed and thus not selected upon. Under the adult recombination statistics an unproductive zero insertion sequence is likely to be paired with a productive sequence with many insertions, and thus we would not expect to see a similar enrichment for unproductive sequences if a general peripheral selective advantage was causing the enrichment. Data source: Emerson *et al.* [2].

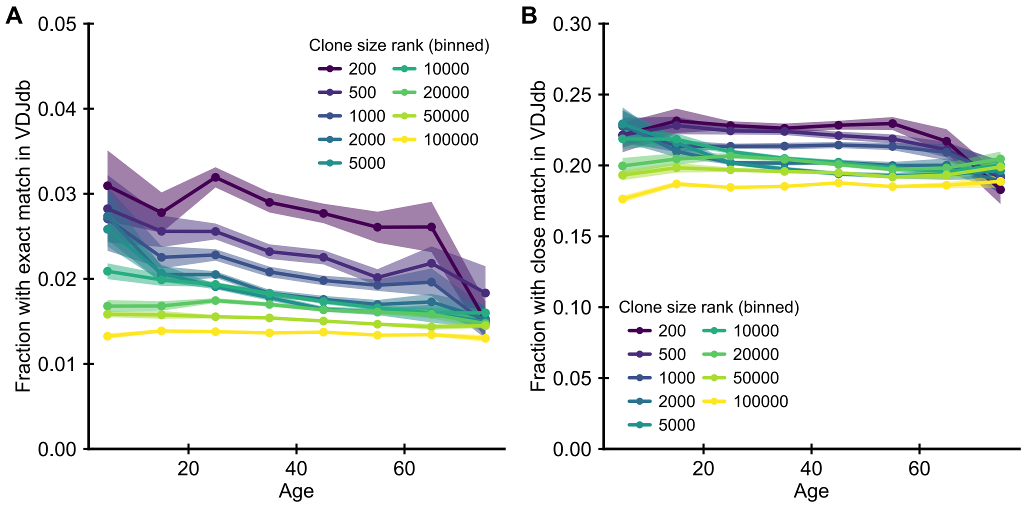


FIG. S6: **Large clones are enriched in clones with known specificity.** (A) Fraction of clones with TCRs that have exact matches in the VDJdb [6] of known antigen specificities. (B) Fraction of clones with close matches (defined as nearest neighbor sequences in a Levenshtein distance sense, i.e. sequences with a single amino acid substitution, insertion or deletion). T cells known to be specific to particular antigens are enriched among the most abundant clones. However, there is little change in this enrichment as a function of age. Data source: Emerson *et al.* [2].

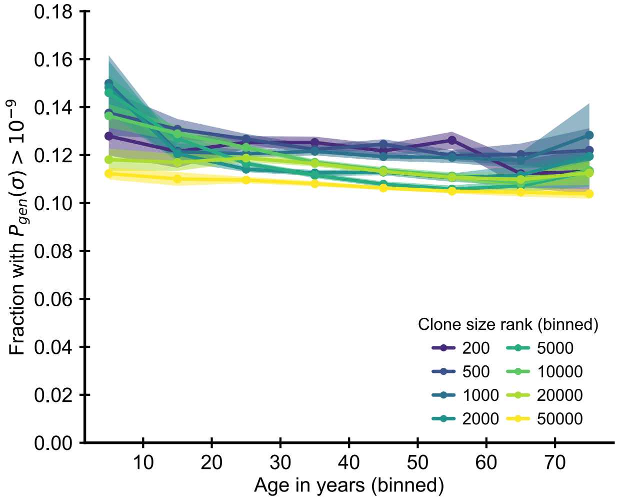


FIG. S7: **Large clones are enriched in clones that are likely to be convergently recombined.** Fraction of clones with TCR sequences $\sigma$ with a probability of generation $P_{gen}(\sigma)$ higher than $10^{-9}$. The probability of generation was calculated based on the nucleotide sequence using a probabilistic model of recombination with default parameters for human TCR sequences [7]. To remove confounding by the early expansionary dynamics we excluded zero insertion clones as most of these clones also have high probability of generation. We find that clones with high $P_{gen}$ are moderately more likely to be large. In comparison to the zero insertion clones, there is little change in their enrichment as a function of age. Data source: Emerson *et al.* [2].

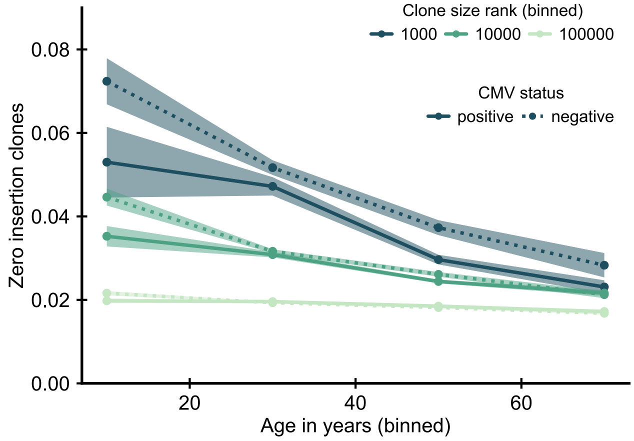


FIG. S8: **Influence of CMV infection status on enrichment of zero insertion clones.** Data source: Emerson *et al.* [2].

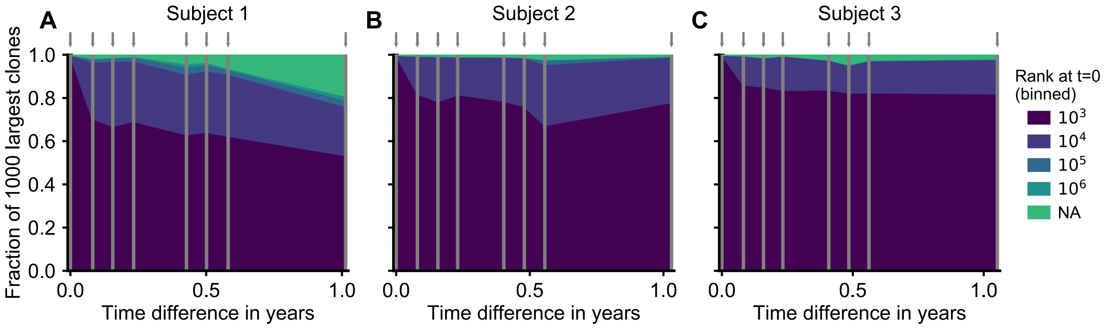


FIG. S9: **Provenance of large T cell clones in a longitudinal study of T cell repertoire dynamics.** Longitudinal analysis of the origin of the 1000 largest clones at each time point (indicated by arrows) in three healthy adults over a one year time frame. For each clone we determined whether it was also sampled at the earliest time point, and if so at what clone size. The plot displays the fraction of clones that fall within a specific clone size rank bin at the first time point. At all times a majority of clones was already large initially. A small fraction was not detected at all at the first time point (ND) likely representing recently expanded clones. (Supplement to Fig. 3D which corresponds to panel C.) Data source: Chu *et al.* [8].

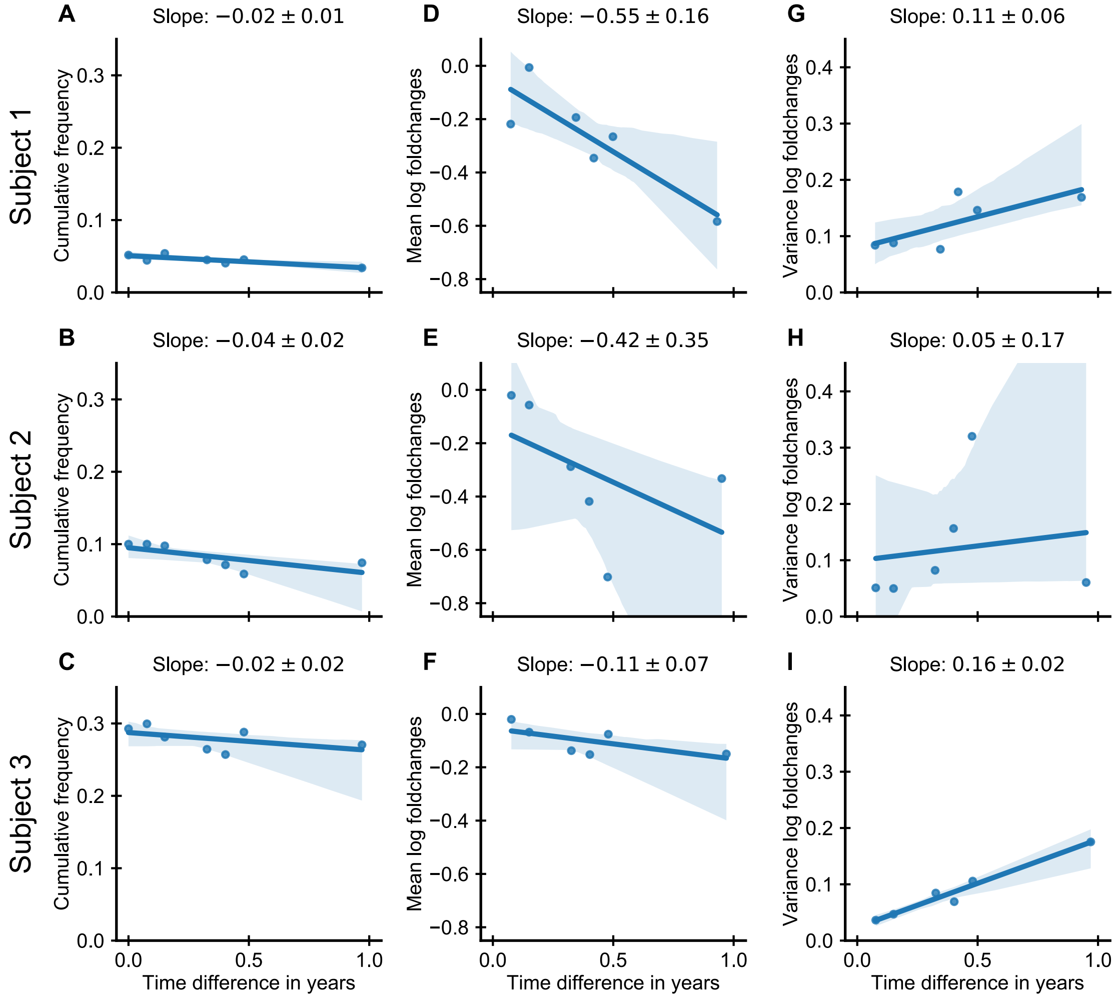


FIG. S10: **Dynamics of large persistent T cell clones in a longitudinal study of T cell repertoire dynamics.** Dynamics of the 250 largest clones from second time point onwards excluding those not sampled at the first time point. (A-C) Fraction of the repertoire represented by these clones (sum of their normalized clone sizes); (D-F) mean and (G-I) variance of the log-foldchanges of their normalized clone sizes relative to time point 2. (Supplement to Fig. 3E which corresponds to panel I.) Data source: Chu *et al.* [8].

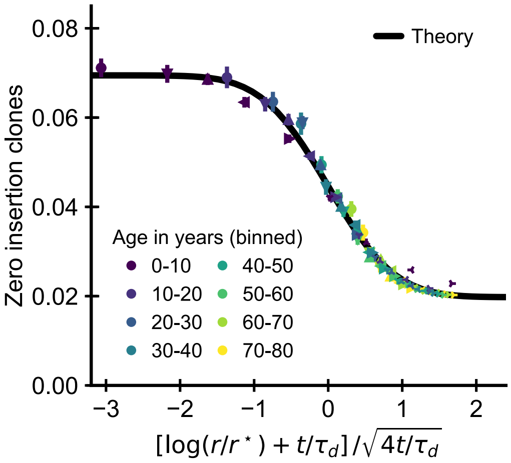


FIG. S11: **Data collapse by parameter rescaling for the simulated cohort.** Same data as in Fig. 3F displayed with a rescaled x-axis using fitted parameters $\tau_d = 10.2 \pm 0.4\,\text{years}, r^\star = 1.19 \pm 0.08 \cdot 10^4$. The data collapses onto a sigmoidal function predicted by theory (SI Text Eq. 51) with fitted $p_{0,-} = 0.0695 \pm 0.0012$, $p_{0,+} = 0.0198 \pm 0.0003$ (black line).

### B. EXTENDED METHODS

#### 1. Data sources

For all studies we used data, which was preprocessed as described in the original study. This data is publicly available from `https://doi.org/10.5281/zenodo.826447` (Britanova cohort), `https://doi.org/10.21417/B7001Z` (Emerson cohort), `https://doi.org/10.21417/PL2018JI` (data from [4]), and `https://doi.org/10.21417/B7J01X` (longitudinal study).

The Britanova cohort comprises 71 individuals spanning ages $6 - 103$ years, as well as 8 cord blood samples. The Emerson cohort spans ages $1 - 74$ years and consists of a training and validation set of 666 and 120 individuals, respectively. From the training set we excluded 111 samples with missing age information and 62 samples with a conflicting data format. We used only samples from the training set to analyze how the scaling law of repertoire organization changes with age (Fig. 1C,E). For the zero insertion enrichment analyses (Fig. 3B,C) we combined both the training and validation set together with separately published repertoire sequencing data from 8 elderly individuals [4] generated using the same experimental pipeline (immunoSEQ, Adaptive Biotechnologies, Seattle) to achieve the broadest possible coverage of all age groups.

The longitudinal study by Chu *et al.* [8] performed repertoire sequencing of peripheral blood from three healthy female volunteers (using the immunoSEQ pipeline) over 8 time points spanning a $\sim$1 year time frame. One individual in the study was in mid-adulthood (24-45 years, Subject 3 in the original study), while two were in early adulthood (18-24 years, Subject 1 and 2 in the original study). In the main text Fig. 3D we display data from the older individual as we expect dynamics of large clones to be masked less by measurement noise as the large clones increase in relative abundance with age.

All studies from which we analyzed data sequenced the locus coding for the TCR CDR3 $\beta$-chain only, and we thus define clones as collections of cells sharing the same CDR3 $\beta$-chain. Clone sizes are defined as the number of distinct unique molecular identifiers (UMIs) sequenced (Britanova cohort), or based on sequencing reads (Emerson cohort). The definition of a clone solely based on the CDR3 $\beta$-chain neglects convergent recombination of the most easily produced receptors with different CDR3 $\alpha$-chains, but we expect convergent recombination to be sufficiently rare overall for this distinction not to qualitatively affect clone size distributions.

We also used flow cytometry data on the fraction of naive cells from Britanova *et al.* [9] and from Shearer *et al.* [10].

#### 2. Data analysis

**Fitting power-law exponents.** We estimate the power-law exponent from sampled clone sizes $\{C_i\}$, $i = 1, \dots, M$, which exceed a minimal size $C_{min}$ by numerically maximizing the log-likelihood of the data [11],

$$\mathcal{L} = -M \ln \zeta(1+\alpha, C_{min}) - (1+\alpha) \sum_{i=1}^{M} \ln C_i, \tag{1}$$

where $\zeta(x,k)$ is the incomplete Riemann zeta function. We use $C_{min} = 16$ for both cohorts, which provides a balance between minimizing bias of the estimated exponents induced by subsampling while not overly increasing the variance of the estimator by excluding most of the data (see Fig. S13).

**Fitting the zero insertion profiles.** To fit the zero insertion fractions to the theory prediction (Eq. 51) we determine the values for $r^\star$ and $\tau_d$ by a weighted least squares fit. We set $r$ and $t$ to the mid-value of each bin for the data. We weight each value by its empirical standard error with an additional model specification error that we set to a fixed value of $2 \cdot 10^{-3}$. To demonstrate the feasibility of the parameter inference we reinferred the parameters from the simulated data and recovered those used as parameter values for the simulation. We also fitted the values of $p_{0,-}$ and $p_{0,+}$, but we note that they are not used in the rescaling and are only needed to display the theoretical curve (Eq. 51).

**Normalization of clone sizes.** Variations in sampling depth can confound comparisons of clone sizes (SI Text C). Intuitively, if we sample more cells overall we also expect to sample proportionally more cells belonging to each given clone. This suggests to use the frequency with which cells are sampled from a given clone as a more robust measure, which can be empirically estimated by normalizing each clone size by the total sample size. We further normalize clone sizes by the fraction of memory T cells found in people of different ages to account for the increase in memory cell fraction in peripheral blood with age (SI Text D). Together these two normalization steps lead to a large degree of data collapse as compared to unnormalized clone sizes.

**Regression analyses.** We determine 95% confidence intervals on regression lines by bootstrapping using case resampling [12].

#### 3. Simulation procedures

**Repertoire formation.** To simulate the model efficiently at large scales we use a mean-field competition approximation (SI Text F 2). We verified the validity of the mean-field assumption by comparing them to full stochastic simulations of the coupled birth-death-immigration equations, which we simulated using the Gillespie algorithm [13] (Fig. S16). In the mean field approximation the proliferation rate is time-dependent, which requires a specific procedure for sampling event times. The time interval until the next event depends on the total rate for all possible processes $\lambda(t) = \theta + b(t) + d$. To sample an interval of time $\Delta t$ between two events from an inhomogeneous Poisson process of rate $\lambda(t)$ one can sample from a Poisson process with a rate function $\lambda^\star(t)$ fulfilling the majoration condition $\lambda^\star \geq \lambda(t) \, \forall t$ and then reject a proposed time interval $\Delta t^\star$ with a probability of $1 - \lambda(t + \Delta t^\star)/\lambda^\star(t + \Delta t^\star)$ [14]. The thinned set of event times follows the statistics of the Poisson process with rate $\lambda(t)$. Here, because competition is increasing with time, $\lambda(t)$ decreases monotonically. Therefore, the homogeneous Poisson process with a constant rate function $\lambda^\star(t) = \lambda(t_0)$, satisfies the majoration condition. Using this thinning technique we are able to efficiently sample the next event time while accounting for the time-dependence of the proliferation rate.

**Simulated cohort.** As empirical evidence shows that the tail of the clone size distribution is almost exclusively driven by cells with memory phenotype (SI Text D), we focused on the clone size dynamics within the memory compartment. We assumed that the recruitment size for memory cells is independent of the prior naive cell dynamics, and we thus did not explicitly model the clone size dynamics within the naive compartment. Within the memory compartment we modeled clone size dynamics under the combined effect of early deterministic expansions during repertoire formation and fluctuating clonal growth rates according to Eq. 6. Given the large sizes of memory clones we expect demographic stochasticity to be negligible relative to clone size variability introduced by fluctuating selection. For tractability we thus ignored demographic fluctuations, which allowed us to combine the continuum solution to the deterministic clonal growth (Eq. 22) with the stochastic propagator for the fluctuating dynamics (Eq. 37) to efficiently simulate the dynamics. To study the enrichment of zero insertion clones in silico we assigned newly recruited memory clones as having zero insertions with a probability equal to the fraction $p_0(t)$ of zero insertion clones within the naive compartment. We assumed $p_0(t) = p_{0,-}$ before TdT expression turn-on at time $t^\dagger$ and $p_0(t) = p_{0,-} t/t^\dagger + p_{0,+}(1-t/t^\dagger)$ for $t > t^\dagger$, where $t/t^\dagger$ is the fraction of naive clones produced since the switch to the adult recombination statistics. Taken together, these simplifications lead to the following direct sampling scheme:

- Sample the age $T$ of an individual uniformly from the range $[0, 80]$ years.
- Set the number of clones equal to $\theta T$ (rounded to the nearest integer), where $\theta$ is the rate of recruitment of new clones to the memory compartment.
- For each clone determine its recruitment time $t_i$ by drawing uniformly from the range $[0, T]$.
- Assign each clone as having zero insertions with a probability

$$p_0(t) = \begin{cases} p_{0,-} & t < t^\dagger \\ p_{0,-} t/t^\dagger + p_{0,+}(1 - t/t^\dagger) & \text{otherwise} \end{cases}$$

- Sample the size $C_i(T)$ of each clone as follows (Eqs. 22 and 37),

$$C_i = \exp(x_i), \quad x_i \sim \mathcal{N}\left(-d(T - t_i) + \frac{1}{1+\gamma} \log\left(\frac{e^{dT} - 1}{e^{dt_i} - 1}\right) - \sigma^2 (T - t_i), 2\sigma^2 (T - t_i)\right), \tag{2}$$

  where $d, \gamma, \sigma^2$ are model parameters and $y \sim N(\mu, \sigma^2)$ indicates x being drawn from a normal distribution of mean $\mu$ and variance $\sigma^2$.

- Finally to mimick the experimental sampling depth of $N_{sample}$ reads we determine sampled clone sizes $\tilde{C}_i$ by Poisson sampling,

$$\tilde{C}_i \sim \mathsf{Pois}(N_{sample} \cdot C_i / N), \quad \text{with } N = \sum_i C_i, \tag{3}$$

  where $x \sim \mathsf{Pois}(\lambda)$ indicates x being drawn from a Poisson distribution of parameter $\lambda$.

#### 4. Parameter choices

In the following we provide a summary of parameter choices we used to simulate repertoire dynamics along with additional motivation.

Lifetimes of several years and several months have been measured by deuterium labelling for naive and memory T cells, respectively [15, 16]. Clonal turnover can be substantially slower than cellular turnover when proliferation balances most death (SI Text E 2). This has been shown to be the case for the maintenance of naive cells in human [17], where the aging-associated decline of the fraction of T cells with T cell receptor excision circles (TRECs) suggests $\gamma \sim 0.1$. Similarly, memory T cell numbers decline much more slowly overall than suggested by the deuterium labelling literature, which is thought to be driven by homeostatic proliferation in the absence of reinfection [18]. For example, T cell memory has been observed to decline with half-lifes of $8-15$ years by following titers after small pox vaccination [19]. Additionally, the relatively short average lifetime of memory T cells likely masks substantial heterogeneity with a subset of more long-lived cells also contributing to the slower long-term decline of memory cells [20]. Another line of direct evidence for long clonal persistence has come from two studies of identical twins [21, 22], which have shown an excess sharing of identical clones decades after *in utero* blood exchange in monochorionic twins.

To simulate repertoire formation (Fig. 2B) we used the following set of parameters:

| parameter | explanation | value |
|---|---|---|
| $d$ | death rate | 0.2/year |
| $\gamma$ | recruitment-to-proliferation ratio | 0.1 |
| $\theta$ | recruitment rate | $10^6$/year |
| $C_0$ | recruitment size | 1 |

We note that under mean-field competition the rate of recruitment $\theta$ only determines the overall number of clones, but does not influence the dynamics of an individual clone and thus the normalized clonal ranks. We thus used a rate smaller than suggested by estimates of thymic output, but importantly large enough to sufficiently sample from the tail of the clone size distribution. The dynamics can furthermore be non-dimensionalized by choosing units where the death rate is one. Therefore the qualitative nature of the results presented in Fig. 2B only depends on $\gamma$, in a way that is shown in Fig. 2D.

To study the enrichment of zero insertion clones in a simulated cohort (Fig. 3E) we used the same recruitment-to-proliferation ratio and death rate as in the previous simulation of repertoire formation. To determine the absolute number of large clones that have zero insertions in these simulations the choice of the recruitment rate $\theta$ is important. Based on order-of-magnitude estimates of the clonal diversity of the memory compartment [23, 24] we chose a value of $\theta = 10^5$/year. Additionally, we chose a fraction of zero insertion clones within the early naive compartment of $p_{0,-} = 0.07$ (roughly equal to their overall fraction in cord blood [21]) and in the late naive compartment equal to $p_{0,+} = 0.02$ (roughly equal to their overall fraction in adult blood). Finally, we used $t^\dagger = 0.05$ years for the time of the recombination switch, which together with the choice of $\theta$ produces $\sim 10^4$ excess zero insertion clones recruited during repertoire formation in line with the enrichment data in the $< 10$ years age group (Fig. 3B). All parameters are summarized in the following table:

| parameter | explanation | value |
|---|---|---|
| $\sigma^2$ | magnitude of clone size fluctuations | 0.08/year |
| $d$ | death rate | 0.2/year |
| $\gamma$ | recruitment-to-proliferation ratio | 0.1 |
| $\theta$ | recruitment rate | $10^5$/year |
| $p_{0,-}$ | Zero insertion fraction early in life | 0.07 |
| $p_{0,+}$ | Adult zero insertion fraction | 0.02 |
| $t^\dagger$ | Time of recombination statistics switch | 0.05 years |
| $N_{sample}$ | simulated sample size | $5 \cdot 10^5$ |

### C. SUBSAMPLING SCALING

Only a small fraction of the $\sim 10^{12}$ T cells in the human body are sampled by repertoire sequencing. What effect does subsampling have on the clone size distribution? In the following we discuss how subsampling affects the distribution of sampled clone sizes and we discuss analysis techniques for robust inferences and data visualization despite variations in sampling depth.

#### 1. Inference of scaling exponent

Given a clone of size $C$ in the repertoire, the number of reads from that clone $\tilde{C}$ follows a distribution $P(\tilde{C}|C)$. The form of $P(\tilde{C}|C)$ depends on the sampling process. To build intuition let us consider the simplest case, in which every cell is sampled independently with a probability $\eta$, the subsampling fraction. Then the sampling distribution is binomial

$$P(\tilde{C}|C) = \binom{C}{\tilde{C}} \eta^{\tilde{C}} (1-\eta)^{C-\tilde{C}}. \tag{4}$$

The mean of this distribution is

$$\langle \tilde{C} \rangle = \eta C, \tag{5}$$

which implies that sampled clone sizes are on average smaller by a factor $\eta$ than the actual clone size. In the practically relevant limit where the sampling fraction is small, $\eta \ll 1$, we can further simplify and assume that the counts from the large clones follow a Poisson distribution. In the Poisson limit the sampled clone size varies around its mean value with a coefficient of variation that scales as an inverse of the square root of the mean sampled count,

$$c_v = \frac{\sqrt{\langle \left( \tilde{C} - \langle \tilde{C} \rangle \right)^2 \rangle}}{\langle \tilde{C} \rangle} = \frac{1}{\sqrt{\eta C}}. \tag{6}$$

Importantly, the stochastic sampling introduces a subsampling scale, $\tilde{C} = \eta C \sim 1$, at the clone size $C = 1/\eta$, from which on average we expect a single sampled cell. Due to the existence of this scale subsampling breaks scale-invariance: even if $P(C)$ follows a perfect power law, the distribution of sampled counts

$$P(\tilde{C}) = \sum_C P(C) P(\tilde{C}|C) \tag{7}$$

deviates from power-law scaling close to $\tilde{C} = 1$. This intuition can be made rigorous using a generating function formalism [25]: for example for $P(C) = C^{-2}/\zeta(2)$ one obtains for $\tilde{C} > 1$

$$P(\tilde{C}) \sim \frac{1}{\tilde{C}(\tilde{C}-1)}. \tag{8}$$

As expected the scaling with an exponent $-2$ is recovered asymptotically, but subsampling leads to a deviation from scaling when $\tilde{C}$ is close to 1.
The deviation from scaling due to subsampling leads to biases in naive estimates of the scaling exponent. How can we determine a power-law exponent in a way that is robust to subsampling? When the sampling distribution is known or can be inferred from replicate sequencing the exponent can be inferred using maximum likelihood estimation of a model with an underlying power law distribution of clone sizes convolved with the sampling probability [26]. Here, we propose a simpler approach that does not require precise knowledge of the sampling process. We exploit the fact that the deviations from scaling vanish asymptotically for large $\tilde{C}$ (Eq. 8), by excluding small clones below some minimal size $C_{min}$ from the fitting. The power-law exponent is expected to converge as we increase $C_{min}$, which we confirm using simulated data (Fig. S12, blue line). We can also consider more realistic models for the sampling process that account for overdispersion, i.e. their coefficient of variation exceeds the minimal value of one set by Poisson sampling. Mechanistically, such overdispersion arises for a number of reasons, most importantly because in practice we are not actually directly counting cells: in the DNA-based sequencing pipeline every cell can give rise to multiple sequencing reads due to the polymerase chain reaction amplification step, and in the mRNA-based sequencing pipeline despite the addition of unique molecular identifiers several of them can originate from different mRNA molecules from the same cell. As long as the number of reads from each cell is independently and identically distributed the law of large numbers ensures that the relative frequencies of large clones converge. We thus expect that the trimming method of fitting only to counts greater than $C_{min}$ also works for overdispersed sampling. We test the trimming method on simulated data, in which the sampling follows a negative binomial distribution with mean $\mu$ and variance $\mu + a\mu^2$ (which reduces to Poisson sampling for $a = 0$). We find that trimming allows a correct estimate of $\alpha$ (Fig. S12, orange and green line). Applying the same method to the empirical data we find that the fitted exponents also depend on $C_{min}$ (Fig. S13). In practice, we chose $C_{min} = 16$ to balance a trade-off between minimizing bias and variance, which increases as more of the data is excluded from the fit (Fig. S13 insets).

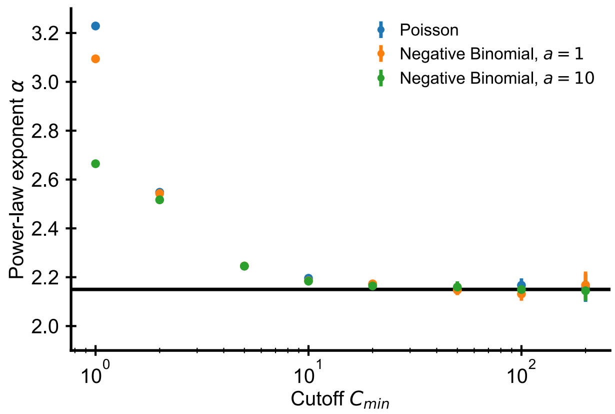


FIG. S12: **Estimated power-law exponents converge to correct value using trimming method.** Fitted exponent as a function of the cutoff choice in simulated data (errorbars $\pm 2 \cdot$ SE over 50 independent draws). The fitted exponent changes drastically for small $C_{min}$ before levelling off indicating deviations from true power-law scaling at the smallest clone sizes. Such a deviation is expected due to subsampling despite the true power-law scaling in the underlying distribution (see text). Simulations: $10^7$ clones were drawn from a discrete power-law distribution with $\alpha = 2.15$. A sample of size $5 \cdot 10^5$ cells was then drawn from the underlying power law based on a Poisson (blue dots) or negative binomial sampling (orange and green dots show two choices of the overdispersion coefficient $a$).

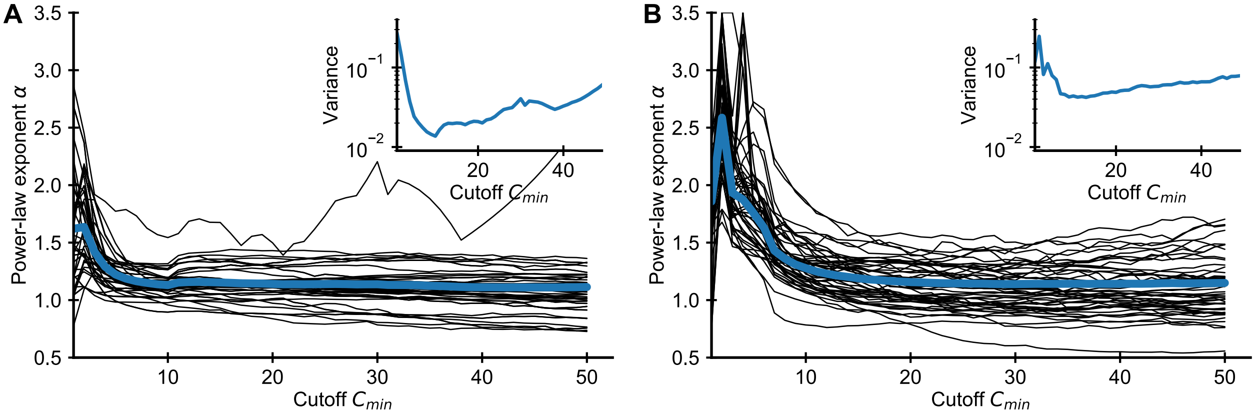


FIG. S13: **Influence of choice of $C_{min}$ on fitted power-law exponent for empirical data.** Fitted exponent as a function of the cutoff choice (black lines: 50 random repertoires, blue line: mean) in the (A) Britanova *et al.* and (B) Emerson *et al.* datasets. The fitted exponent changes drastically for small $C_{min}$ before levelling off indicating deviations from true power-law scaling at the smallest clone sizes, similarly to those seen in simulated data (Fig. S12). To alleviate the bias induced by finite sampling we choose a cutoff value $C_{min}$, for which the power-law exponent estimates have levelled off. For large $C_{min}$ the variance of fitted exponent increases as more and more data is excluded from the fit (A, B inset), which sets a practical upper bound for choosing $C_{min}$.

#### 2. Graphical display of subsampled distributions

The intuition we have built about how subsampling affects clone size distributions can help us choose an appropriate method for displaying subsampled data (Fig. S14). Which graphical representation of the clone size distribution minimizes the influence of variations in sampling depth?
The shift of the mean clone size (Eq. 5) suggests that we should normalize sampled clone sizes by the sampling fraction $\eta$, as has been noted elsewhere [27]. While experimentally we do not know the sampling fraction, we can instead simply divide the clone sizes by the total sample size (Fig. S14C,D). This normalization is particularly intuitive as it corresponds to using the relative frequencies of cells in different clones. While the absolute number of cells in a

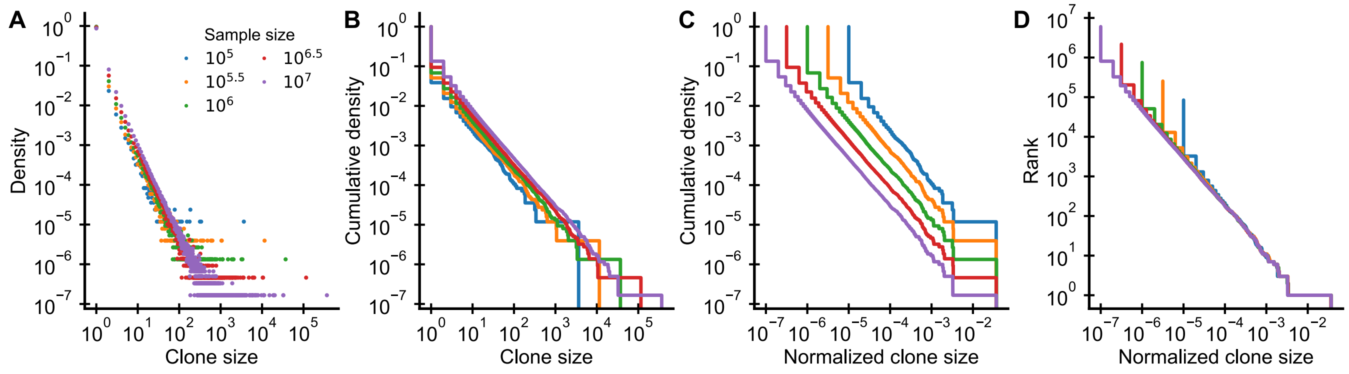


FIG. S14: **Graphical display of subsampled power-law distributions.** (A-D) show various ways of displaying clone size distributions obtained by subsampling an underlying clone size distribution consisting of $10^8$ clones drawn according to $P(C) \sim C^{-2.2}$ to various sampling depths. (A) The empirical probability density function of clone sizes, (B) its cumulative density, as well as (C) the cumulative density of normalized clone sizes are not invariant under changes of the sampling depth. Only the tail behavior of relative frequencies of finding cells from large clones is reproducibly captured, which makes rank-frequency plots (displays of unnormalized cumulative distributions of normalized clone sizes) the method of choice for collapsing clone size distributions at various sampling depths.

large clone increases with more sampling, the fraction of all sampled cells that are part of a particular clone remains constant on average.

Plots of the cumulative distribution of clone sizes make it easier to visually assess the tail behavior of the distribution (Fig. S14B) than plots of the probability density (Fig. S14A). However, even after normalizing clone sizes by the sample size there remains a very visible shift between the cumulative distributions at different sampling depths (Fig. S14C). This shift arises because the implicit normalization by the total number of unique clones makes the sampled cumulative distribution depend heavily on sampling depth. As sampling increases so does the total number of unique clones that will be sequenced. This suggests that we might do better by simply omitting the normalization. Ranking clones by their normalized size yields precisely such an unnormalized cumulative distribution. Taken together, by both scaling clone sizes by the sample size and resisting the temptation to normalize the ranks, we can collapse distributions sampled at different depths (Fig. S14D).

### D. RELATION BETWEEN CLONE SIZE AND CELLULAR PHENOTYPES

In both cohorts all T cells from peripheral blood were sequenced irrespective of their phenotypes. Antigenic challenges drive large clonal expansions and we thus expect clones with effector or memory cells to be larger than naive clones all else being equal [28, 29]. This has generally been confirmed by TCR repertoire sequencing studies [30], but there have also been some reports [21, 24] of expanded naive clones with similar sizes to the largest memory clones. Given this unclear picture from the literature we analyzed the relative contribution of naive and memory cells to clones of different sizes.

Overall, we might expect that naive clones dominate the clone size distribution at the smallest sizes. To test this idea we compared sequencing and flow cytometry data from the Britanova cohort and founnd that the fraction of naive cells in different individuals explains a remarkably high 88% of variability in the number of clones sequenced only once after subsampling all repertoires to the same size (Fig. S15A). To further determine how cells from clones of different sizes partition phenotypically we analyzed data from a study in which T cells were sequenced both in unsorted blood as well as after sorting into naive and memory cells [8]. We find that the sizes of large clones follow the same scaling in unsorted blood and in the memory compartment (Fig. S15B). Within the naive compartment most clones are small, in particular when excluding clones from which cells are also found in the memory compartment (Fig. S15B, red line). We note from the plot that all of the largest 200 clones in unsorted blood have memory phenotype cells, and less than one percent of the top 1000 clones are not found within the memory compartment. This rules out that the enrichment of zero insertion clones among the most abundant clones found in Fig. 3 is driven by naive clones as has been suggested in a previous study [21]. The relative frequency of a clone within the memory compartment is larger by a constant fold-factor (Fig. S15C), likely reflecting an increased relative frequency of the large clones when excluding naive cells from the denominator.

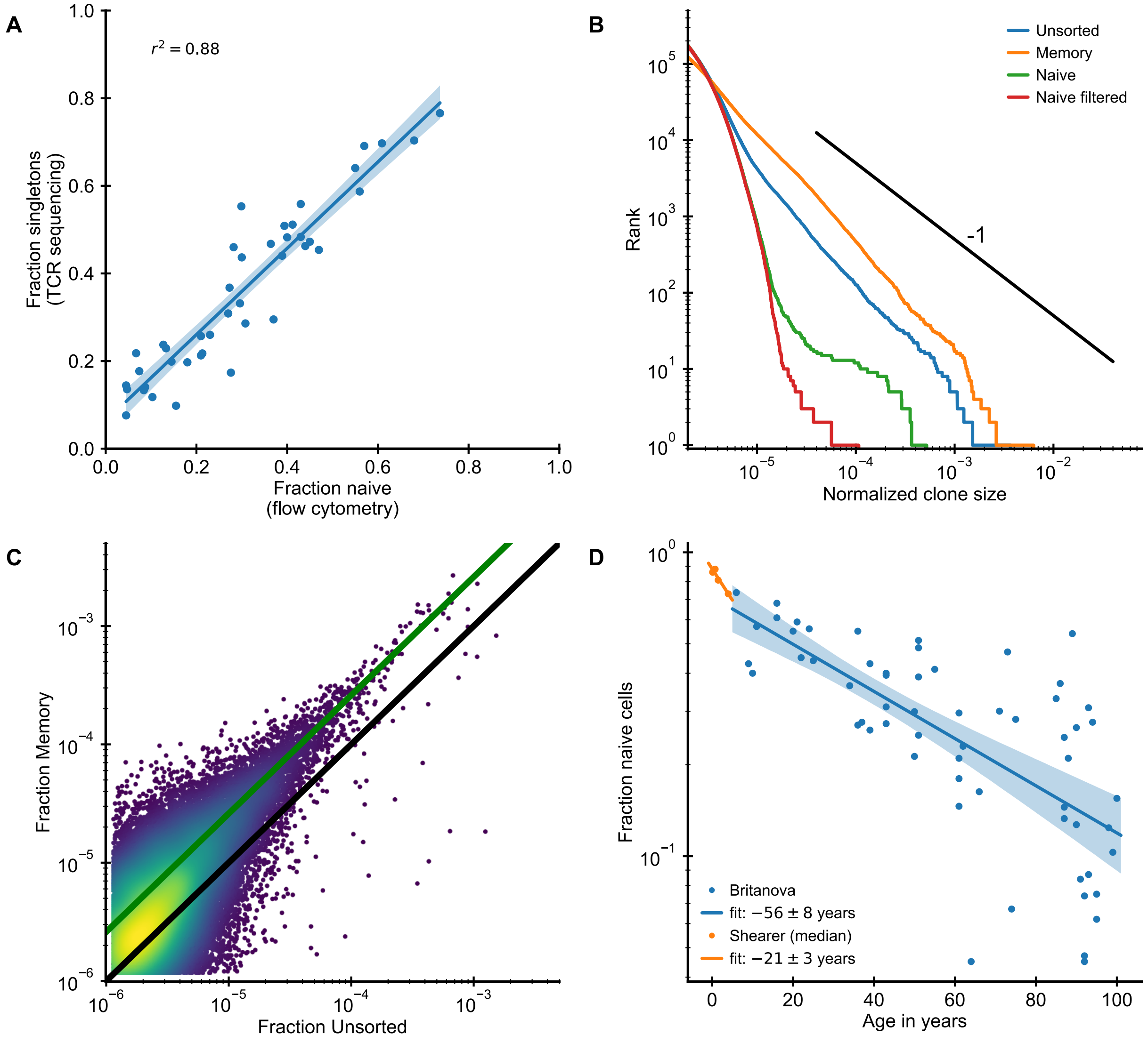


FIG. S15: **The large clones in unsorted peripheral blood are predominantly of memory phenotype.** (A) The naive cell fraction as determined by flow cytometry and the fraction of singletons are closely correlated in the Britanova cohort. To diminish the influence of sampling depth variations we computationally subsampled all repertoires to an equal sample size of $5 \cdot 10^5$ counts. (B,C) Analysis of unsorted (TCR sequencing from all peripheral blood mononuclear cells), memory ($CD3^+$, $CD45RO^+$), and naive ($CD3^+$, $CD45RA^+$) blood samples from the same individual (Data source: [8]). (A) Clone size distributions in the different T cell compartments. Filtering naive clones that are also found in the memory compartment removes most large naive clones. (B) Frequency of large clones in the memory sample is shifted upwards relative to their frequency within the unsorted sample. Color represents logarithm of local kernel density estimate in regions with overplotting. The solid lines are guides to the eye (black line represents equal frequency, green line 2.6-fold higher frequency in the memory compartment). (D) Fraction of naive cells decreases with age (Data source: [1]) starting in early infancy (Data source: [10]). Legend shows fitted time constant of exponential decay (± SE).

To correct for the decrease of naive cells with age (Fig. S15D) [1, 10] we normalize clonal frequencies in unsorted peripheral blood by the mean fraction of memory cells expected at different ages fit to the flow cytometry data. We find that this normalization collapses the tails of empirical clone size distributions (Fig. S2C,F).

### E. MODELING NEUTRAL REPERTOIRE DYNAMICS

In the following we review results on the neutral dynamics of clone sizes in which the continuous recruitment of new clones is balanced by a net negative growth of already established clones $b < d$. These models have a long history in ecology [31], and have also been proposed as null models in the context of T cell dynamics previously [32–34]. While the results can be found in the literature their inclusion serves to introduce a parametrization highlighting the recruitment-to-proliferation ratio $\gamma$ as a key quantity governing clonal dynamics.

#### 1. Steady state clone size distribution

At steady state the probability distribution $P(C)$ of clone sizes $C$ needs to fulfill the balance condition

$$bCP(C) = d(C+1)P(C+1), \tag{9}$$

for all $C > C_0$ which yields

$$P(C) \propto \frac{1}{C}\left(\frac{b}{d}\right)^C = \frac{1}{C}\exp\left(-C\log(d/b)\right). \tag{10}$$

This distribution is characterized by power-law scaling with an exponent of 1 for small clone sizes, and, importantly, has an exponential cutoff at $C^\star = 1/\log(d/b)$. In contradiction with this model experiments point towards power-law scaling with an exponent ∼2 (Note that $P(C) \sim C^{-\alpha-1}$ when rank $\sim C^{-\alpha}$). Additionally the size of large clones seen experimentally is incompatible with the predicted exponential cutoff as we discuss below.
The total repertoire size follows the following continuum equation

$$\frac{\mathrm{d}N}{\mathrm{d}t} = (b-d)N + \theta C_0, \tag{11}$$

such that at steady-state, $\frac{\mathrm{d}N}{\mathrm{d}t} = 0$, the repertoire has a total size

$$N_\infty = \frac{\theta C_0}{d-b}. \tag{12}$$

For a more interpretable alternative parametrization we introduce the recruitment-to-proliferation ratio for the maintenance of cells at steady state

$$\gamma = \frac{\theta C_0}{bN_\infty} = \frac{d}{b} - 1. \tag{13}$$

Using this relation to rewrite Eq. 10 we obtain

$$P(C) \propto \frac{1}{C}\exp\left(-C\log(1+\gamma)\right), \tag{14}$$

implying a cutoff clone size of $C^\star = 1/\log(1+\gamma)$. The largest clones represent on the order of one percent of the repertoire, which assuming independent sampling from the underlying repertoire would correspond to $\sim 10^{10}$ cells in the complete repertoire. For small $\gamma$ we can expand $C^\star \approx 1/\gamma$, so in order to have a cutoff clone size $C^\star$ of this order of magnitude one would need to have an unreasonably small $\gamma \sim 10^{-10}$.

#### 2. Relaxation time scale

Over what timescale do transiently expanded clones disappear? The time scale $\tau_c = \frac{1}{d-b}$ for deterministic clonal decay can be much larger than the lifespan $1/d$ of a single cell when birth and death are closely balanced. Rewriting the birth rate in terms of $\gamma$ and $d$ we obtain

$$\tau_c = \frac{1+\gamma}{d\gamma}, \tag{15}$$

demonstrating that for $\gamma \ll 1$ clonal dynamics is a factor of $1/\gamma$ slower than cellular dynamics.

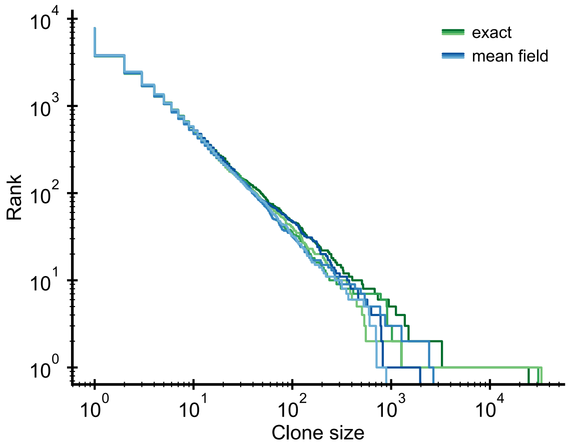


FIG. S16: **Validation of the mean-field approximation.** Comparison of full stochastic simulations and simulations using mean-field competition. Parameter: $b_0 = 2 \cdot 10^4$/year, $d = 0.2$/year, $\theta = 2 \cdot 10^3$/year (implying $\gamma = 0.1$), simulation length 5 years.

### F. MODELING REPERTOIRE FORMATION

#### 1. Mechanistic motivation for the competition function

We consider a population of $N$ T cells that proliferate at a rate proportional to the concentration $S$ of a set of stimuli (stimulatory cytokines), $b \propto S$. We assume that the cytokines are produced by other cells at some fixed rate $p$ and degraded at a basal rate $q$. We further assume that competition between T cells is mediated by their consumption of cytokines. The dynamics of $S$ is then described by

$$\frac{\mathrm{d}S}{\mathrm{d}t} = p - qS - kSN, \tag{16}$$

where $-kSN$ is a mass action term describing how T cells lower cytokine levels. Assuming a separation of timescales in which cytokine concentrations change quickly we obtain the quasi steady state approximation

$$S = \frac{p}{q + kN}. \tag{17}$$

When the consumption term dominates relative to basal decay, $kN \gg q$, we obtain $b \propto S \propto 1/N$.

#### 2. Mean-field competition approximation

We simplify the full stochastic model (Eqs. 2-4) using a mean-field approximation for the competition, which decouples the dynamics of individual clones while retaining the full stochasticity on the clonal level. This approximation replaces the dependence of the proliferation rate on $N$ by a dependence on its continuum theory average given by Eq. 20. We exactly simulated a system of reduced size to validate the mean-field approximation (see Sec. B 3). The distributions of the exact and mean-field simulations agree to within stochasticity (Fig. S16), with the exception of the largest clone, which is larger in the exact simulations as has been discussed elsewhere [35].

#### 3. Continuum theory of clonal growth

To obtain insight into why the model produces power-law scaling we present a simple continuum theory of early clonal dynamics. We approximate the clone size dynamics of the $i$-th clone $C_i$ as

$$\frac{\mathrm{d}C_i}{\mathrm{d}t} = \left(\frac{b_0}{N(t)} - d\right) C_i, \tag{18}$$

with $C_i(t_i) = C_0$ at the time of recruitment $t_i$. The total repertoire size $N = \sum_i C_i$ evolves according to

$$\frac{\mathrm{d}N}{\mathrm{d}t} = b_0 - dN + \theta C_0, \tag{19}$$

whose solution is given by

$$N(t) = (b_0 + \theta C_0)\left(1 - e^{-d\,t}\right)/d. \tag{20}$$

For times large compared to $1/d$ the total repertoire size given in Eq. 20 reaches a steady-state,

$$N_\infty = (b_0 + \theta C_0)/d, \tag{21}$$

because competition for proliferation signals acts as a homeostatic regulator. By combining Eq. 20 and Eq. 18 we derive the clonal growth law

$$C_i(t) = C_0 \left(\frac{e^{dt} - 1}{e^{dt_i} - 1}\right)^{1/(1+\gamma)} e^{-d(t-t_i)}, \tag{22}$$

where $\gamma$ as in SI Text E is the recruitment-to-proliferation ratio which in this model is given by $\gamma = \theta C_0/b_0$. To simplify we expand the growth law at leading order for small times, $t_i < t \ll 1/d$, to obtain

$$C_i(t) = C_0 \left(\frac{t}{t_i}\right)^{1/(1+\gamma)}. \tag{23}$$

This expression can also be derived directly by noting that early repertoire growth is linear $N(t) \approx (b_0 + \theta C_0)t$, and that the early dynamics is dominated by proliferation and not death such that

$$\frac{\mathrm{d}C_i}{\mathrm{d}t} = \frac{1}{(1+\gamma)t} C_i, \tag{24}$$

which is solved by Eq. 23. Given the constant recruitment of new clones the distribution of the $t_i$'s is uniform, which with Eq. 23 implies a clone size distribution

$$P(C) = P(t_i(C)) \left|\frac{\mathrm{d}t_i}{\mathrm{d}C}\right| \propto C^{-2-\gamma} \tag{25}$$

that follows power-law scaling with an adjustable exponent that depends on $\gamma$. Note that the exponent for $P(C)$ differs by one from the exponent for the rank [11], which is a complementary cumulative distribution, and thus $\alpha = 1 + \gamma$.

#### 4. Steady-state distribution

To derive the power-law scaling we have expanded the total repertoire size for small times (or death rates). How does the clone size distribution change later in life? At large times the division rate $b_0/N(t)$ falls below the constant death rate $d$ as the steady-state repertoire size $N_\infty$ is approached following Eq. 20. In this model this happens at a time $t^\star \simeq \log(1 + 1/\gamma)/d$, after which the large clones experience a deterministic force towards extinction. For times $t \gg t^\star$ the model effectively reduces to the neutral birth-death dynamics considered in SI Text E. (The growth rate fluctuations produced by variations of the total population size around steady state asymptotically vanish for large $N_\infty$.) We thus expect the steady-state clone size distribution to be equivalent to that of the neutral model (Eq. 14). Indeed this distribution accurately describes the distribution of small clones in old age (Fig. 2B). The neutral distribution is not compatible with data as discussed before. However, the timescale over which large early founded clones vanish is long (SI Text. E 2) such that a tail of large clones resulting from the early growth dynamics can be maintained much beyond $t^\star$ until $t \gg \tau_c$.

#### 5. Relaxations of model assumptions

For tractability and interpretability we have kept the model presented in the main text deliberately simple. Here, we explore how a saturation of the proliferation rate, competition for specific resources, or variations in the recruitment size modify clone size distributions.

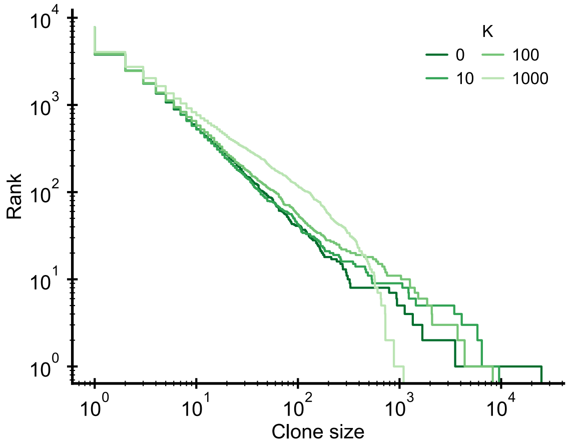


FIG. S17: **Saturation of proliferation rate.** Influence of a saturation of the proliferation rate, $b = b_0/(K+N)$, on the clone size distribution. The saturation induces a change of the scaling behavior at the largest clone sizes. Parameter: $b_0 = 2{\cdot}10^4$/year, $d = 0.2$/year, $\theta = 2 \cdot 10^3$/year (implying $\gamma = 0.1$), simulation length 5 years.

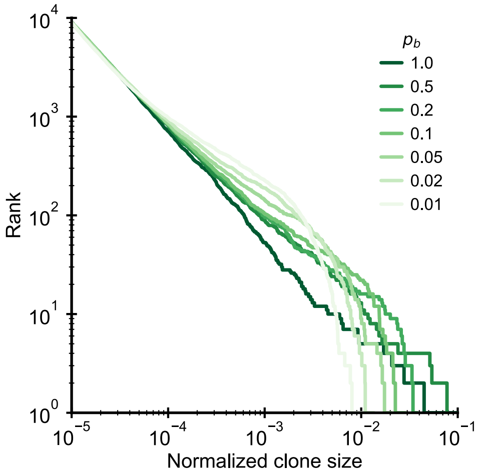


FIG. S18: **Competition for specific resources.** Clone size distributions in a simulated model where clones compete for specific antigens to which they bind with a probability $p_b$. Parameter: $b_0 = 10^4$/year, $\theta = 10^3$/year (implying $\gamma = 0.1$), $N_a = 1000$, $d = 0$, simulation length 10 years.

**Saturation of proliferation rate.** Cellular growth is not arbitrarily fast, which is not accounted for in the simple model in which cells proliferate very rapidly early in life. To understand how such a saturation effect influences clone size distributions we introduce an upper limit on birth rate that limits proliferation in the absence of competition. Following [36] we set the clonal birth rate to $b(t) = b_0/(K + N)$ for some constant $K$, which sets the repertoire size below which competition is negligible. Given this choice the birth rate remains limited to a value $b_0/K$ even in the absence of any competitors. Increasing $K$ leads to deviation in the scaling of the largest clones (Fig. S17), but the same scaling remains at intermediate clone sizes. In the model early clonal growth is exponential until the total repertoire has reached size $N(t) \sim K$, which explains the different distribution of the largest clones. However, the number of clones that are recruited during this phase grows only logarithmically with $K$ due to the exponential increase in total repertoire size.

**Competition for specific resources.** T cells respond to stimuli from peptide-MHC complexes, which could also act as limiting resources. T cells then compete only with those cells specific to the same antigens in contrast to the global competition considered previously. To assess how assumptions about the mechanisms of competition influence

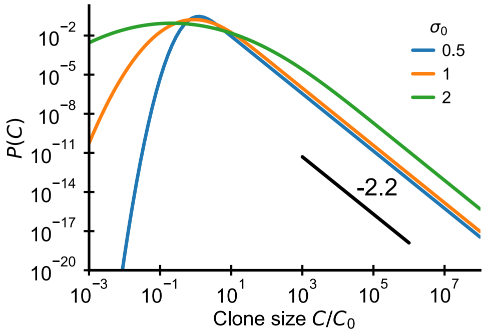


FIG. S19: **Variation of recruitment size.** Clone size distributions resulting from a variable recruitment size and repertoire growth (Eq. 28). The black line shows a power law with a slope of -2.2 for visual comparison. Parameter: $\gamma = 0.2$

our results we simulated the repertoire formation process using a classical description of competition for antigens [37–39]. We consider a fixed number of antigens $N_a$ and encode the specificity of the $M$ clones in a matrix $K$ of size $M \times N_a$, where $K_{ij} = 1$ if clone $i$ recognizes the antigen $j$ and $K_{ij} = 0$ otherwise. We draw the entries of $K$ independently with a fixed binding probability $p_b$. We assume that the proliferation rate of a cell of the $i$-th clone is proportional to the amount of antigenic stimulation:

$$b_i = \frac{b_0}{N_a} \sum_{j=1}^{N_a} K_{ij} F_j \tag{26}$$

where the availability of antigen $j$ is given by

$$F_j = \frac{1}{1 + \sum_i K_{ij} C_i}. \tag{27}$$

The normalization of Eq. 26 ensures that total proliferation is comparable to a global resource model with the same parameters independent of $N_a$. For computational tractability we simulated the clone size dynamics without taking into account demographic stochasticity in proliferation and death of cells. While more specific competition (smaller $p_b$) leads to a deviation in the distribution of the largest clones, we find that clone size distributions are heavy tailed independently of the choice of $p_b$ and all display the same scaling at intermediate clone sizes (Fig. S18).

**Variations of the recruitment size.** The numbers of cells $C_0$ that are recruited might also be variable. In particular, this will be the case when we think about the memory compartment, in which $C_0$ represents the number of cells from a clone recruited into memory following infection. To understand how such variations modify the dynamics of repertoire formation we derive an analytical prediction in the case where the distribution of recruitment sizes, $P(C_0)$, is lognormal. Given a lognormal distribution with parameters $\mu_0$ and $\sigma_0$ the mean introduction size is given by $\overline{C_0} = e^{\mu_0 + \sigma_0^2/2}$. To keep the mean introduction size constant while changing the variability of clone sizes, we use a parametrization in terms of $\overline{C_0}$ and $\sigma_0$ and set $\mu_0 = \log(\overline{C_0}) - \sigma_0^2/2$. To determine the clone size distribution resulting from early repertoire growth we integrate the continuum theory prediction, $P(C/C_0) \propto (C/C_0)^{-2-\gamma}$ over the distribution of $C_0$:

$$\begin{aligned} P(C) &\propto \int_0^C dC_0 \left(C/\overline{C_0}\right)^{-2-\gamma} \frac{1}{\sigma_0 C_0/\overline{C_0}\sqrt{2\pi}} e^{-\left(\log\left(C_0/\overline{C_0}\right)+\sigma_0^2/2\right)^2/(2\sigma_0^2)} \\ &= \left(C/\overline{C_0}\right)^{-2-\gamma} \cdot e^{\frac{1}{2}(\gamma+2)(\gamma+1)\sigma_0^2} \frac{1}{2} \operatorname{erfc}\left(\frac{-2\log(C/\overline{C_0}) + (3+2\gamma)\sigma_0^2}{2\sqrt{2}\sigma_0}\right). \end{aligned} \tag{28}$$

The complementary error function erfc$(x)$ saturates for $x \ll -1$ and thus the distribution follows the same power-law $P(C) \sim C^{-2-\gamma}$ for large clones, $\log C/\overline{C_0} \gg \sigma_0 \left(\sqrt{2} + \frac{3+2\gamma}{2}\sigma_0\right)$, while it deviates for smaller clones within the range of recruitment sizes (Fig. S19).

#### 6. Relation to mechanisms generating power laws in other growth processes

The origin of power law scaling during repertoire formation is reminiscent of a class of stochastic processes widely studied in the literature as a mechanism underlying power-law distributions found in diverse contexts [40–42], which has been rediscovered multiple times since the pioneering work of Yule on speciation [40]. Common to these processes is that the distribution of types at a given point is the result of a balance between the growth of existing types and the addition of new types. The different models depending on their context differ in (i) the growth rate $r(t)$ of the number of units of each already existing type and (ii) the rate function $\theta(t)$ at which new types are introduced. They all share the same basic mathematical mechanism that produces a power law distribution of types as we review below. The three maybe most well-known instances of this class of processes are the following:

- the Yule model of speciation [40], in which (i) species within a genus speciate at some constant rate, and (ii) new genus is created at a rate proportional to the number of already existing genera.
- the Luria-Delbrück model of bacterial population genetics during exponential growth [41], in which (i) each cell divides at a constant rate, and (ii) new alleles arise through random mutation at a constant rate per cell division.
- the Barabási-Albert (BA) model of network growth [42], in which at every time step (ii) a new node is added, and is (i) linked to $m$ already existing nodes with a probability proportional to the number links that a chosen node already has.

Despite differing in their assumptions about the functional form of the growth and innovation rates we show in the following that these different models all share a common mathematical basis. To provide a common terminology we will use the language of urn models and refer to different types as urns and to the different number of units of each types as balls in each urn. In an attempt to unify the different models we develop a continuum theory for these growth-innovation processes. To do so we rescale time to

$$\tau = \int_0^t \theta(t')\mathrm{d}t', \tag{29}$$

such that new urns are added at unit rate, $\theta(\tau) = 1$. The number of balls in each urn then grows according to

$$\frac{\mathrm{d}C_i}{\mathrm{d}\tau} = \frac{\mathrm{d}C_i}{\mathrm{d}t}\frac{\mathrm{d}t}{\mathrm{d}\tau} = \frac{r(t(\tau))}{\theta(t(\tau))}C_i =: \zeta(\tau)C_i. \tag{30}$$

The key to the power-law scaling in all these models is the existence of a regime in which

$$\zeta(\tau) = \frac{1}{\alpha\tau}, \tag{31}$$

i.e. the growth rate scales inversely with rescaled time with a proportionality factor $1/\alpha$. Eq. 31 has the same form as Eq. 24 that we derived for our model of repertoire formation. Thus following the derivation of Eq. 25 within SI Text F 3 we obtain a subexponential growth of balls in already existing urns, which leads to a power law scaling of the distribution of balls per urn,

$$P(C) \propto C^{-\alpha-1}, \tag{32}$$

with an adjustable exponent that depends on $\alpha$.

Before deriving how Eq. 31 arises in specific contexts let us first remark on a general consequence of this form of effective growth law: The total number of balls added to all existing urns per rescaled time unit is constant. To derive this let us assume that each new urn is populated by $C_0$ balls, then the total number of balls $N(t) = \sum_i C_i$ grows according to

$$\frac{\mathrm{d}N}{\mathrm{d}\tau} = \frac{1}{\alpha\tau}N + C_0, \tag{33}$$

which is solved by

$$N(\tau) = \frac{C_0\alpha}{\alpha - 1}\tau. \tag{34}$$

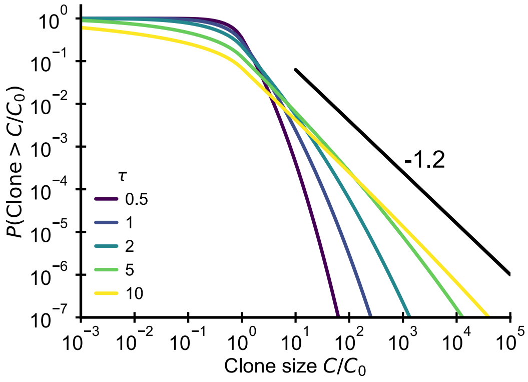


FIG. S20: **Fluctuating fitness model out-of-steady state.** Analytical predictions for the clone size distributions in a geometric Brownian motion fluctuating fitness model (Integral of Eq. 42) as a function of effective age $\tau = T\sigma^2$. The black line shows the asymptotic prediction for the steady-state scaling. Parameter: $\alpha = 1.2$

Multiplying by the growth rate Eq. 31 yields a constant,

$$N(\tau)\zeta(\tau) = \frac{C_0}{\alpha - 1}, \tag{35}$$

thus showing the equivalency between the assumed growth rate depency on rescaled time and the constancy of how many balls are added per rescaled time unit.

In the Yule process we have $r(t) = r$ and recruitment is proportional to the number of genera, $\theta(t) = sG(t)$, which grow at a (generally different) rate $s$, $G(t) = G_0 e^{st}$. By integration we obtain $\tau(t) = G_0\left(e^{st} - 1\right)$, which leads to $\zeta(\tau) = \frac{r}{s\tau + sG_0}$. Thus $\zeta(\tau) \approx \frac{r}{s\tau}$ when the number of newly created genera exceeds the initial number $\tau \gg G_0$. The exponent of the power-law is determined by the ratio of the growth of genera and species, $\alpha = s/r$.

In the Luria-Delbrück model we have $r(t) = r$; recruitment is proportional to the total population size $\theta(t) = \mu r N(t)$, where $\mu$ is the mutation probability per replication and where $N(t) = N_0 e^{rt}$. By integration we obtain $\tau(t) = \mu N_0\left(e^{rt} - 1\right)$, which leads to $\zeta(\tau) = \frac{1}{\tau + \mu N_0}$. Thus $\zeta(\tau) = \frac{1}{\tau}$ when $\tau \gg \mu N_0$. In contrast to Yule's model the power-law exponent is fixed at $\alpha = 1$, because the same growth process governs the increase in $\theta(t)$ and in cell numbers.

In the Barabasi-Albert model the introduction rate $\theta(t) = 1$ is constant, but $r(t)$ decreases with time. The $m$ newly added links attach preferentially to those nodes that already have a large degree. The growth rate $r(t) = m/N(t)$ of a node thus decreases proportionally to the total degree $N = 2mt$ of all present nodes. We have $r(t) = 1/(2t)$, which implies $\zeta(\tau) = \frac{1}{2\tau}$ and $\alpha = 2$.

### G. MODELING LONG-TERM REPERTOIRE DYNAMICS WITH FLUCTUATING CLONAL GROWTH RATES

#### 1. Slow convergence to steady-state scaling

Multiplicative stochastic processes are a classical generative mechanisms for heavy-tailed distributions [43–45]. In the context of lymphocyte dynamics this mechanism has first been proposed by Desponds *et al.* [32], who argued that fluctuations in antigen availability can lead to multiplicative stochastic dynamics producing power-law scaling at steady state. Here, we expand on this ealier work by analyzing a simple fluctuating fitness model out-of-steady-state. Our analytical results show that the emergence of scaling can be slow when the fluctuation amplitude is small.

We opted to treat proliferation rate fluctuations as temporally uncorrelated for computational tractability (Eq. 6). Correlations in proliferation rate fluctuations are clearly an important feature of short term dynamics – e.g. to describe the quick expansion and contraction during and following acute infection over a timescales of days and weeks, respectively [29]. However, given finite correlation times we expect to be able to capture dynamics over the long timescales which we are interested in here, with uncorrelated noise with an effective net fluctuation strength that averages over the short-term dynamics.

In this limit clone sizes follow a geometric Brownian motion, i.e. $x = \log C/C_0$ follows the Langevin equation

$$\frac{\mathrm{d}x_i}{\mathrm{d}t} = f_0 + \sqrt{2}\sigma\eta_i, \tag{36}$$

with initial condition $x(t_i) = 0$, where $\sigma$ sets the fluctuation strength and where $\langle \eta_i(t)\eta_j(t')\rangle = \delta_{ij}\delta(t-t')$. A negative mean fitness $f_0 < 0$ balances the recruitment of new clones and the net expansion induced by the fluctuating term. In general, we might want to include also demographic noise and the extinction of clones as an absorbing boundary condition [32], but here for simplicity we will neglect those effects. Eq. 36 is a diffusion equation for the logarithmic clone size $x$ and has the well-known Green's function

$$G(x, y, t) = \frac{1}{\sqrt{4\pi\sigma^2 t}} e^{-\frac{(x-y-f_0 t)^2}{4\sigma^2 t}}, \tag{37}$$

which describes how the distribution spreads out from an initial $\delta$-distribution centered at size $y$. The clone size distribution at time $T$ is given by

$$P(x, T) = \int_0^T \mathrm{d}t P(t) G(x, 0, t), \tag{38}$$

where $t$ is the clonal age. For a constant immigration rate $t$ is uniformly distributed and we obtain by integration

$$P(x, T) = \frac{e^{\frac{f_0 x(1-\theta(x))}{\sigma^2}} \operatorname{erfc}\left(\frac{|x|-f_0 T}{\sqrt{4T\sigma^2}}\right) - e^{\frac{f_0 x\theta(x)}{\sigma^2}} \operatorname{erfc}\left(\frac{|x|+f_0 T}{\sqrt{4T\sigma^2}}\right)}{2 f_0 T}, \tag{39}$$

where $\theta(x)$ is the Heaviside step function, $\theta(x) = 0$ for $x < 0$ and $\theta(x) = 1$ otherwise. For large $T$ and $x > 0$ this reduces to

$$P(x) \to e^{\frac{f_0 x}{\sigma^2}} / (-f_0 T), \tag{40}$$

which implies

$$P(C) \sim C^{-(1+\alpha)} \quad \text{with}\, \alpha = -f_0/\sigma^2, \tag{41}$$

recovering the steady-state result from [32].
Setting $f_0 = -\alpha\sigma^2$ and rescaling age as $\tau = T\sigma^2$, we can rewrite the finite time solution as

$$P(x, \tau) = \frac{e^{-\alpha x\theta(x)} \operatorname{erfc}\left(\frac{|x|+\tau\alpha}{\sqrt{4\tau}}\right) - e^{-\alpha x(1-\theta(x))} \operatorname{erfc}\left(\frac{|x|-\alpha\tau}{\sqrt{4\tau}}\right)}{2\alpha\tau}. \tag{42}$$

Plotting the cumulative distribution of clone sizes at different effective ages (Fig. S20) we observe that the convergence of clone size distributions is slow when $\sigma^2$ is small. Based on estimates for the fluctuation strength from longitudinal data (Fig. 3D) we would expect significant deviations from the steady state power-law scaling that persist into adulthood. Thus this mechanism alone is unable to account for the observed power-law scaling in data.

#### 2. A note on the scaling exponent

A minimal requirement for the existence of a steady state is $f_0 < 0$ ensuring that clones eventually die to balance the recruitment of new clones. This condition still allows such multiplicative processes to produce power-laws with arbitrary exponents as noted before [32]. Here, we propose that the parameters should fulfill a stronger condition. In particular, it seems reasonable to require that the large clones do not deterministically take up a larger fraction of the overall repertoire, or equivalently that their expected change in clone size should not exceed one. The mean of the lognormal distribution of clone size change is given by $e^{f_0+\sigma^2}$, and thus we find the stronger condition

$$-f_0 < \sigma^2. \tag{43}$$

Importantly, it follows that exponents in the vicinity of $\alpha = -1$ arise without fine-tuning as long as the timescale of expected net clonal decay is large compared to the diffusion timescale.
Another perspective on the parameterization is provided by noting that the Langevin equation for $C$ (not $x = \log C$) in the Stratonovic convention includes an extra drift term $-\sigma^2$, to keep $\langle \Delta C\rangle$ independent of the choice of $\sigma$. Alternatively, in the Ito convention the extra drift term arises by Ito's lemma when transforming the equation from $C$ to $x$

#### 3. Predictions for longitudinal fluctuations in clone sizes

To quantify longitudinal fluctuations we calculate the mean and variance of log-clonesize changes with respect to a reference time $t_0$. From the model we according to Eq. 37 expect

$$\langle x(t) - x(t_0) \rangle = f_0 t \tag{44}$$

$$\langle (x(t) - x(t_0) - \langle x(t) - x(t_0) \rangle)^2 \rangle = 2\sigma^2 t. \tag{45}$$

The variance of log-clonesize changes in empirical data involves an additional term $\sigma_S^2$ accounting for sample-to-sample variability. This term is expected not to depend on the time difference, and we can thus determine $\sigma^2$ by linear regression with an intercept that captures the sampling variability $\sigma_S^2$ (Fig. 4B).
We note that a similar approach has been independently proposed in unpublished work by Ferri and advisors [46].

#### 4. Relaxation of the zero insertion distribution

Here, we solve for the relaxation dynamics of the zero insertion distribution in a simplified setting. Throughout we use log clone sizes $x = \log C$ for notational convenience. We posit that at time 0 the power-law distribution $P(x, 0) = \alpha e^{-\alpha x}$ is already established and we further assume that the $r^\star$ largest clones have zero insertion probability $p_{0,-}$ and all smaller or later added clones have probability $p_{0,+}$. Then the probability that a clone of a given size $x$ has zero insertions is given by

$$P_0(x, t) = \Delta p_0 f_{early}(x, t) + p_{0,+} \tag{46}$$

where $\Delta p_0 = p_{0,-} - p_{0,+}$ and $f_{early}(x, t)$ is the fraction of clones of size $x$ and time $t$ that derive from the $r^\star$ largest clones at time 0.
In the following we determine an analytical formula for $f_{early}(x, t)$ under the assumption that the dynamics leaves the distribution unchanged $P(x, t) = P(x, 0)$. We then have

$$f_{early}(x, t) = \frac{\int_{x_{min}}^{\infty} \mathrm{d}y \, e^{-\alpha y} G(x, y, t)}{e^{-\alpha x}}, \tag{47}$$

where $G(x, y, t)$ as before is the Green's function of the fluctuating proliferation rate dynamics and $x_{min}$ is defined such that the total number of clones times $P(x > x_{min})$ equals $r^\star$. By integration one obtains

$$f_{early}(x, t) = \frac{1}{2} e^{\alpha t (f_0 + \alpha \sigma^2)} \operatorname{erfc}\left( \frac{x_{min} - x + t(f_0 + 2\alpha\sigma^2)}{\sqrt{4\sigma^2 t}} \right), \tag{48}$$

which after setting $f_0 = -\alpha\sigma^2$ reduces to

$$f_{early}(x, t) = \frac{1}{2} \operatorname{erfc}\left( \frac{x_{min} - x + \alpha\sigma^2 t}{\sqrt{4\sigma^2 t}} \right). \tag{49}$$

To convert clone size into ranks, we note that rank $\sim e^{-\alpha x}$ and thus $x_{min} - x \sim \frac{1}{\alpha} \log\left(\frac{r}{r^\star}\right)$. In combination with Eqs. 49 and 46 we thus obtain

$$P_0(r, t) = \frac{\Delta p_0}{2} \operatorname{erfc}\left( \frac{\frac{1}{\alpha} \log(r/r^\star) + \alpha\sigma^2 t}{\sqrt{4\sigma^2 t}} \right) + p_{0,+}. \tag{50}$$

Defining a characteristic timescale for the diffusive dynamics as $\tau_d = 1/(\alpha\sigma)^2$ we can simplify this expression to

$$P_0(r, t) = \frac{\Delta p_0}{2} \operatorname{erfc}\left( \frac{\log(r/r^\star) + t/\tau_d}{2\sqrt{t/\tau_d}} \right) + p_{0,+}. \tag{51}$$

[1] O. V. Britanova, M. Shugay, E. M. Merzlyak, D. B. Staroverov, E. V. Putintseva, M. A. Turchaninova, I. Z. Mamedov, M. V. Pogorelyy, D. A. Bolotin, M. Izraelson, et al., The Journal of Immunology **196**, 5005 (2016).

[2] R. O. Emerson, W. S. DeWitt, M. Vignali, J. Gravley, J. K. Hu, E. J. Osborne, C. Desmarais, M. Klinger, C. S. Carlson, J. A. Hansen, et al., Nature Genetics pp. 1–10 (2017).
[3] A. W. Sylwester, B. L. Mitchell, J. B. Edgar, C. Taormina, C. Pelte, F. Ruchti, P. R. Sleath, K. H. Grabstein, N. A. Hosken, F. Kern, et al., Journal of Experimental Medicine **202**, 673 (2005).
[4] P. Lindau, R. Mukherjee, M. V. Gutschow, M. Vignali, E. H. Warren, S. R. Riddell, K. W. Makar, C. J. Turtle, and H. S. Robins, The Journal of Immunology **202**, 476 (2019).
[5] S. L. Klein and K. L. Flanagan, Nature Reviews Immunology **16** (2016).
[6] M. Shugay, D. V. Bagaev, I. V. Zvyagin, R. M. Vroomans, J. C. Crawford, G. Dolton, E. A. Komech, A. L. Sycheva, A. E. Koneva, E. S. Egorov, et al., Nucleic Acids Research pp. 1–9 (2017).
[7] Z. Sethna, Y. Elhanati, C. G. Callan, A. M. Walczak, and T. Mora, Bioinformatics **35**, 2974 (2019).
[8] N. D. Chu, H. S. Bi, R. O. Emerson, A. M. Sherwood, M. E. Birnbaum, H. S. Robins, and E. J. Alm, BMC Immunology **20**, 1 (2019).
[9] O. V. Britanova, D. A. Bolotin, E. A. Bogdanova, M. A. Turchaninova, Y. B. Lebedev, S. Lukyanov, I. Z. Mamedov, E. M. Merzlyak, E. V. Putintseva, D. B. Staroverov, et al., The Journal of Immunology **192**, 2689 (2014).
[10] W. T. Shearer, H. M. Rosenblatt, R. S. Gelman, R. Oymopito, S. Plaeger, E. R. Stiehm, D. W. Wara, S. D. Douglas, K. Luzuriaga, E. J. McFarland, et al., Journal of Allergy and Clinical Immunology **112**, 973 (2003).
[11] A. Clauset, C. R. Shalizi, and M. E. J. Newman, SIAM review **51**, 661 (2009).
[12] B. Efron and T. Hastie, *Computer Age Statistical Inference* (2016).
[13] W. H. Press, S. a. Teukolsky, W. T. Vetterling, and B. P. Flannery, *Numerical Recipes 3rd Edition: The Art of Scientific Computing*, vol. 1 (Cambridge University Press, 2007).
[14] P. A. Lewis and G. S. Shedler, Naval research logistics quarterly **26**, 403 (1979).
[15] R. J. De Boer and A. S. Perelson, Journal of Theoretical Biology **327**, 45 (2013).
[16] J. A. Borghans, K. Tesselaar, and R. J. de Boer, Immunological Reviews **285**, 233 (2018).
[17] I. den Braber, T. Mugwagwa, N. Vrisekoop, L. Westera, R. Mögling, A. Bregje de Boer, N. Willems, E. H. Schrijver, G. Spierenburg, K. Gaiser, et al., Immunity **36**, 288 (2012).
[18] D. C. Macallan, J. A. Borghans, and B. Asquith, Vaccines **5** (2017).
[19] E. Hammarlund, M. W. Lewis, S. G. Hansen, L. I. Strelow, J. A. Nelson, G. J. Sexton, J. M. Hanifin, and M. K. Slifka, Nature Medicine **9**, 1131 (2003).
[20] R. S. Akondy, M. Fitch, S. Edupuganti, S. Yang, H. T. Kissick, W. Kelvin, G. Alexe, S. Nagar, M. M. Mccausland, H. A. Abdelsamed, et al., Nature **552**, pages362 (2017).
[21] M. V. Pogorelyy, Y. Elhanati, Q. Marcou, A. L. Sycheva, E. A. Komech, V. I. Nazarov, O. V. Britanova, D. M. Chudakov, I. Z. Mamedov, Y. B. Lebedev, et al., PLoS Computational Biology **13**, e1005572 (2017).
[22] H. Tanno, T. M. Gould, J. R. Mcdaniel, W. Cao, Y. Tanno, and R. E. Durrett, Proceedings of the National Academy of Sciences **117** (2020).
[23] H. S. Robins, P. V. Campregher, S. K. Srivastava, A. Wacher, C. J. Turtle, O. Kahsai, S. R. Riddell, E. H. Warren, and C. S. Carlson, Blood **114**, 4099 (2009).
[24] Q. Qi, Y. Liu, Y. Cheng, J. Glanville, D. Zhang, J.-Y. Lee, R. a. Olshen, C. M. Weyand, S. D. Boyd, and J. J. Goronzy, Proceedings of the National Academy of Sciences of the United States of America **111**, 13139 (2014).
[25] M. P. Stumpf, C. Wiuf, and R. M. May, Proceedings of the National Academy of Sciences **102**, 4221 (2005).
[26] M. Puelma Touzel, A. M. Walczak, and T. Mora, arXiv preprint arXiv:1912.08304 (2019).
[27] A. Levina and V. Priesemann, Nature Communications **8** (2017).
[28] D. L. Farber, N. A. Yudanin, and N. P. Restifo, Nature Reviews Immunology **14**, 24 (2014).
[29] A. Mayer, Y. Zhang, A. S. Perelson, and N. S. Wingreen, Proceedings of the National Academy of Sciences **116**, 5914 (2019).
[30] T. Oakes, J. M. Heather, K. Best, R. Byng-Maddick, C. Husovsky, M. Ismail, K. Joshi, G. Maxwell, M. Noursadeghi, N. Riddell, et al., Frontiers in Immunology **8**, 1 (2017).
[31] I. Volkov, J. R. Banavar, S. P. Hubbell, and A. Maritan, Nature pp. 1035–1037 (2003).
[32] J. Desponds, T. Mora, and A. M. Walczak, Proceedings of the National Academy of Sciences **113**, 274 (2016).
[33] J. Desponds, A. Mayer, T. Mora, and A. M. Walczak, arXiv preprint arXiv:1703.00226 (2017).
[34] P. C. D. Greef, T. Oakes, B. Gerritsen, M. Ismail, J. M. Heather, R. Hermsen, B. Chain, R. J. D. Boer, M. James, R. Hermsen, et al., eLife **9**, e49900 (2020).
[35] P. S. Dodds, D. R. Dewhurst, F. F. Hazlehurst, C. M. V. Oort, L. Mitchell, A. J. Reagan, J. R. Williams, and C. M. Danforth, Physical Review E **052301**, 1 (2017).
[36] R. J. De Boer and A. S. Perelson, Journal of Theoretical Biology **175**, 567 (1995).
[37] R. J. De Boer and A. S. Perelson, Journal of Theoretical Biology **169**, 375 (1994).
[38] R. J. De Boer, A. A. Freitas, and A. S. Perelson, Journal of Theoretical Biology **212**, 333 (2001).
[39] A. Mayer, V. Balasubramanian, T. Mora, and A. M. A. Walczak, Proceedings of the National Academy of Sciences **112**, 5950 (2015).
[40] G. U. Yule, Phil. Trans. B **213**, 21 (1924).
[41] S. E. Luria and M. Delbrück, Genetics **28**, 491 (1943).
[42] A. L. Barabási and R. Albert, Science **286**, 509 (1999).
[43] D. Sornette and R. Cont, Journal de Physique **1**, 431 (1997).
[44] X. Gabaix, The Quarterly Journal of Economics **114**, 739 (1999).
[45] M. E. Newman, Contemporary Physics **46**, 323 (2005).

[46] S. Ferri, *Master thesis: Stochastic processes for natural evolutionary dynamics of T-cell repertoires* (2018).
